# Global transcriptional regulators fine-tune the translational and metabolic machinery in *Escherichia coli* under anaerobic fermentation

**DOI:** 10.1101/2020.07.17.209353

**Authors:** Mahesh S. Iyer, Ankita Pal, Sumana Srinivasan, Pramod R. Somvanshi, K.V. Venkatesh

## Abstract

Complex regulatory interactions between genetic and metabolic networks together confer robustness against external and internal perturbations in an organism such as *Escherichia coli*. In balanced exponential growth, this robustness is attributed to cost-effective metabolism by means of efficient resource allocation coordinated by the interplay of global transcriptional regulators with growth-rate dependent machinery. Here, we reappraise the role of global transcriptional regulators FNR, ArcA and IHF, integral to sustaining proteome-efficiency in anaerobic fermentative conditions, fundamental for optimal growth of *E. coli*. We reveal at the transcriptome and metabolome level, that absence of these global regulators ensued a disruption of nitrogen homeostasis, overexpression of otherwise unnecessary or hedging genes and impairment in core bottleneck steps and amino acid metabolism. Notably, our findings emphasize their importance in optimizing the metabolic proteome resources essential for rapid exponential growth. Consequentially, the perturbations in the metabolic proteome as a result of deletion of global regulators unbalances the ribosomal proteome share imposing a high translation program, though at the expense of lowered efficiency. We illustrate that disruption of this inherent trade-off between metabolic and ribosomal proteomic investment eventually culminate to lowered growth rates. Despite no changes in gene expression related to glucose import, our findings elucidate that the accumulations of intracellular metabolites directly modulated by growth rate, negatively impacts the glucose uptake. Our results employing the proteome allocation theory and quantitative experimental measurements, suffices to explain the physiological consequences of altered translational and metabolic efficiency in the cell, driven by the loss of these global regulators.

## Introduction

In gram-negative bacteria, *E. coli*, due to the huge expenditure associated with protein synthesis especially ribosomes, a tight coordination of the proteomic resources with the metabolic needs is fundamental for cell proliferation (1–3). Broadly, this can be explained by the trade-off existing between metabolic flux involved in generating the precursors and amino acids, and protein synthesis flux utilizing the amino acids, that results in balanced exponential growth (4). Such metabolic and translational capacities facilitated by central carbon metabolic proteome and ribosomes, entails bacterial growth to be precisely controlled by the joint effect of global physiological factors (e.g. RNA polymerase, ribosome, metabolites) and transcriptional regulators (4–6). Additionally, resource investments need to be carefully tailored to specific environmental conditions, that enables organism to objectively distribute the proteome share towards energy and biomass synthesis (7).

The global transcriptional regulators, FNR (Fumarate and Nitrate Reductase), ArcA (Aerobic respiration control A) and IHF (Integration host factor) occupy the top-most hierarchy of transcriptional regulatory network (TRN) based explicitly on the number of genes they regulate (8). Genome-wide binding studies for global regulators like FNR and ArcA in varied oxygenation and nutritional conditions are well appreciated (9–12). Conventionally, broad overview of the role of these regulators in sensing metabolic redox state, influence of oxygen on their activity and the regulatory networks governing central carbon metabolic responses have been thoroughly investigated using gene expression, mutational studies and metabolic flux analyses in anaerobic, micro-aerobic and anaerobic-aerobic transition environments (10–20). Similarly, IHF has been well-characterized as a nucleoid protein and for its regulatory effects under diverse nutritional conditions (21–24). However, a fundamental question as to how the transcriptional regulators orchestrate efficient allocation of proteome share towards metabolism and ribosome biogenesis that collectively accounts for biomass synthesis, has not yet been addressed. Such studies employing regulatory effects of FNR, ArcA and IHF, can unravel the underlying molecular mechanisms that lead to metabolic and translational efficiencies in an energetically least favourable anaerobic fermentative condition. Therefore, a detailed investigation of the causality of these regulator deletions on the phenotype of the organism perceived as shifts in growth optimality, will provide a holistic understanding of how the organism initiates quantitative cellular responses to environmental cues.

Here, we elaborate using a systems biology approach (transcriptomics, ^13^C-metabolomics and phenomics), that disruption of FNR, ArcA and IHF independently affect the allocation of global cellular factors (e.g. RNA polymerase, ribosome, metabolites) and causes imbalanced proteome allocation that governs the optimality of exponentially growing cells. We first characterized the transcriptomic responses under glucose fermentative conditions that helped us to identify known and novel signature genes specific to the regulators which overall represented dysregulation of common metabolic pathways. We demonstrated that all these regulators exhibit direct or indirect control over nitrogen sensing mechanism, repress unnecessary genes under anaerobic fermentation and impact the biosynthesis and bottleneck reactions in central metabolic pathway. Collectively, they represent the core metabolic proteome essential for rapid growth and their impairment influences the growth-dependent ribosomal proteome as well as the metabolome of the organism. Overall, utilizing a simple quantitative model (4, 25–28), we revealed that these global regulators fine-tune the metabolic and translational machinery, thereby ensuring sector-specific proteome distribution, conducive for balanced exponential growth. Our work utilizing the in-depth details to derive the macroscopic system-level perspective elucidates the general design principles underlying the importance of global regulators in *E. coli*, that spans more than 30 years of research.

## Results

### Disparate transcriptome profiles of global regulator mutants converge to a common phenomenon of metabolic impairment

First, to address the transcriptomic changes caused by the disruption of global regulators namely FNR, ArcA and IHF in *E. coli* K12 MG1655 under anaerobic glucose fermentative condition, we performed a high coverage RNA-seq in the mid-exponential phase of growth. Preliminary analysis of RNA-Seq data revealed a total of ~314, ~258 and ~580 differentially expressed genes (DEGs) in Δ*fnr*, Δ*arcA* and Δ*ihf* mutants respectively, compared to the wild-type strain (WT) (Fig. S1A). The Δ*fnr* and Δ*arcA* mutant showed a distinct pattern in its DEGs, wherein we observed higher number of downregulated genes (~66%) in Δ*fnr* mutant and higher number of upregulated genes (~67%) in Δ*arcA* mutant (Fig. S1B). This result reiterated their predominant role as transcriptional activator and repressor, respectively (10–12). On the contrary, in *Δihf* mutant, the number of upregulated (53%) and downregulated (47%) genes were comparable with slightly higher median fold-change (FC) in case of downregulated genes (P < 0.05, Mann Whitney test) (Fig. S1B).

We compared our gene expression data with previous studies encompassing gene expression and binding profiles (9–12, 16). We consistently found a significant yet lower percentage of DEGs directly regulated by FNR (Up DEGs: ~11%, P > 0.05; Down DEGs: ~14%, P < 10^-3^), ArcA (Up DEGs: ~30%, P < 10^-4^; Down DEGs: ~29%, P < 10^-4^) and IHF (Up DEGs: ~10%, P < 10^-4^; Down DEGs: ~17%, P < 10^-5^). Moreover, DEGs also showed enrichment for RNA polymerase with either stress-related sigma factor, sigma 38 or nitrogen-related factor, sigma 54 apart from the growth associated sigma factor 70 (Fig. S2A-S2C). This reflected a sub-optimal allocation of growth-rate dependent global machinery as an indirect consequence of loss of these regulators.

Further, we enriched the DEGs for each of the mutants using KEGG pathway hierarchical classification to understand the role of these regulators on key metabolic pathways (Fig. S3-S5). We observed a common enrichment (adj-P value < 0.05) of pathways such as Amino acid metabolism and Transport across all the mutants. Additionally, we observed enrichment of pathways pertaining to TCA cycle and anaplerosis, and alternate carbon metabolism (e.g. lipid and steroid metabolism, glycerol or secondary carbon degradation) in these mutants. Altogether, such enrichments underscore the metabolic impairment prevalent as a result of loss of these global regulators.

### Perturbation in nitrogen homeostasis

We investigated the gene-level changes associated with the enrichment of Amino acid metabolism and Transport in all the mutants (Fig. 1 and Fig. S3-S5). Specifically, we observed upregulation of putrescine (*puuABCE*) and arginine degradation (*astABCDE*) genes in Δ*fnr* and Δ*arcA* mutants. These genes belong to pathways that yield glutamate and ammonia as end products that can independently satisfy *E. coli*’s nitrogen requirement (29, 30). Perhaps, upregulation of these genes indicated a scavenging mechanism to restore possible disruption of the nitrogen balance in the cell. In addition, we observed downregulation of ATP-dependent organic nitrogen transporters namely arginine (*artIJ*) and methionine (*metN*) transporters in Δ*arcA*, and glutamine (*glnHP*) and leucine/phenylalanine (*livHKM*) transporters in Δ*fnr*. These gene responses indicated an impairment in amino acids or nitrogen sources import among these mutants.

**Fig. 1.**
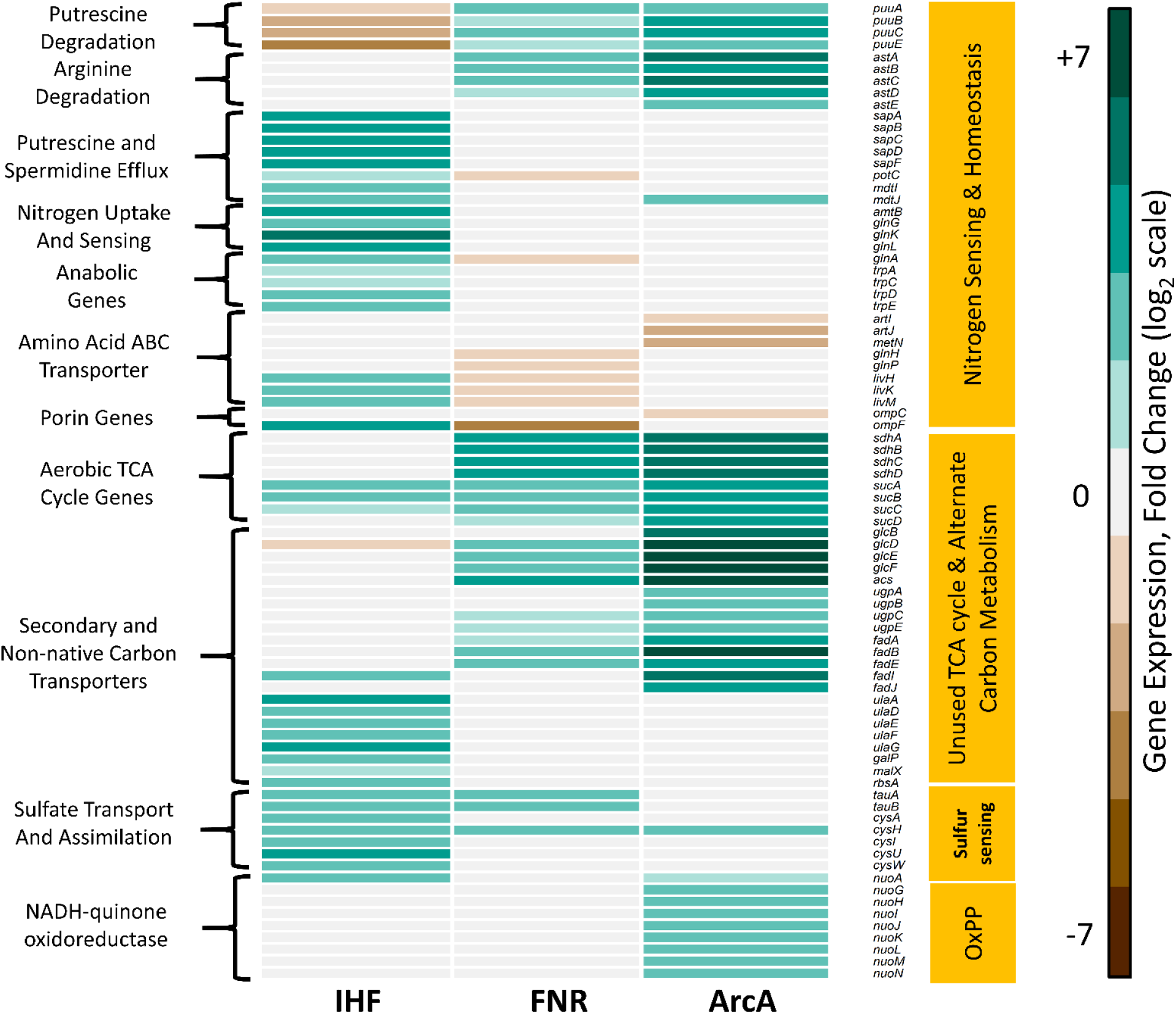
Heatmap of representative DEGs in Δ*ihf*, Δ*fnr* and Δ*arcA* strains, each compared to WT. The figure shows specific genes enriched by KEGG pathway classification (shown in Fig. S3-S5) that were functionally categorised using evidences from Ecocyc database. Broadly, loss of these global transcriptional regulators was found to commonly affect genes involved in nitrogen sensing and homeostasis, unused TCA cycle and alternate carbon metabolism, sulfur sensing and oxidative phosphorylation (OxPP). Gene expression values are obtained from average of two biological replicates (n=2) expressed as log2 Fold change. The downregulated genes are shown in brown and the upregulated genes are shown in cyan.

Similar observation of nitrogen limitation was prevalent in Δ*ihf* (Fig. 1). For instance, upregulated genes enriched for Transporters in Δ*ihf*, code for efflux of polyamines like putrescine (*sapABCDF*) and spermidine (*potC, mdtIJ*), which represent organic nitrogen reservoirs, known to be involved in protein synthesis contributing to cell proliferation (31). Under nitrogen replete conditions, *E. coli* resorts to increasing gene expression of ATP-dependent nitrogen uptake and two-component nitrogen sensing systems, sufficient enough to sustain cell growth (29, 32). Accordingly, we observed upregulation of *amtB and glnKLG* genes in Δ*ihf*. In addition, we observed upregulation of amino acid biosynthesis genes such as glutamine (*glnA*) and tryptophan (*trpACDE*) in Δ*ihf*, in agreement with increased expression of anabolic genes in response to nitrogen limitation (27).

Moreover, we observed changes in outer-membrane porin genes, such as downregulation of *ompF* in Δ*fnr* and *ompC* in Δ*arcA* and upregulation of *ompF* in Δ*ihf* mutants, perceived as hyper-osmotic or hypo-osmotic conditions respectively (Fig. 1). As alteration of these porin genes reflected the osmotic imbalance faced by the cell which in turn has strong associations with nitrogen assimilation, such changes might conform to the defect in nitrogen transport (33–35). Overall, such characteristic expression profiles elucidated an important role of these global regulators in regulating the nitrogen homeostasis and sensing mechanism in *E. coli*.

### De-repression of genes unnecessary in anaerobic fermentation

Our analysis indicated an upregulation of TCA cycle and anaplerosis pathway in Δ*fnr* and Δ*arcA* (Fig. 1 and Fig. S4,S5) that involved genes coding for the aerobic respiratory TCA cycle, in agreement with previous datasets (10–12, 16). Apart from being unnecessary under the anaerobic fermentative conditions, these genes also incur a substantial protein synthesis cost (36). In addition, we observed a significant upregulation of alternate carbon metabolism genes such as glycolate degradation (*glcBDEF*), glycerol degradation (*ugpABCE*) and acetate uptake (*acs*) in Δ*fnr* and Δ*arcA*. This represents a hedging mechanism, wherein a bacterial cell when challenged with fluctuating or unfavorable environmental conditions, modulates its gene expression so as to prepare itself conducive to alternate metabolic phenotypes albeit conforming to the supply and demand conferred on the system (37–39). Furthermore, our DEGs showed significant upregulation of aerobic oxidative phosphorylation (*nuoAGHIJKLMN*) and unused lipid and steroid metabolism *(fadABEIJ)* related pathways in Δ*arcA*.

Similarly, Δ*ihf* showed significant upregulation of alternate carbon metabolism genes responsible for ascorbate (*ulaADEFG*), galactose/maltose (*galP*, *malX*), and ribose (*rbsA*) catabolism (Fig. 1). We further revealed upregulation of genes coding for transporters of sulfur sources (*tauAB* and *cysAUW*) as well as sulfate assimilation (*cysHI*), as a common response to sulfur limitation prevalent in Δ*ihf*. Overall, the upregulation of genes pertaining to unused or hedging proteins under conditions of anaerobic fermentation, can impose a burden on the growth of the organism.

### Regulatory control on amino acid biosynthesis and bottleneck steps in central carbon metabolic pathway

Transcriptionally regulating the first step or final committed step suffices to control the end-product concentrations, be it the amino acid biosynthetic pathway or crucial reactions in central carbon metabolism (36, 40). In order to effectively examine the control on such amino acid biosynthesis and specific bottleneck reactions, we monitored the absolute intracellular concentration of metabolites in the mid-exponential phase using ^13^C-based metabolomics coupled to its corresponding gene expression changes.

Among the precursor metabolites of the glycolytic pathway, a pronounced effect was seen as an increased phosphoenolpyruvate (PEP) accumulation in all the mutants (Fig. 2 and Fig. S6). Presumably, this observation can be attributed to upregulation of *pck* in Δ*ihf* and Δ*arcA* and upregulation of *ppsA* and downregulation of *pykA* in Δ*fnr*, that have been reported to contribute to PEP synthesis during glycolysis (41). Akin to *pykA*, we observed down-regulation of *ackA* gene in Δ*fnr*, both of which contribute to ATP synthesis via substrate level phosphorylation. Also, we observed accumulation of branched chain amino acids derived from pyruvate, namely valine and leucine, in Δ*ihf* and Δ*fnr* mutant (Fig. 2 and Fig. S6). This was in accordance with lowered gene expression of *ilv* observed for both the mutants, thereby indicating a feedback regulation of these amino acids on their biosynthesis. Additionally, in Δ*ihf*, we observed increase in levels of glycerate-3-phosphate (3PG), a precursor for the amino acid glycine and serine, in concert with down-regulation of *gpmA* gene expression (Fig. 2 and Fig. S7). However, only glycine levels were found to be higher in both Δ*ihf* and Δ*fnr*. Taken together, these results suggest that these regulators play a pivotal role in regulating glycolysis, majorly exerted through optimal metabolite levels of the pathway.

**Fig. 2.**
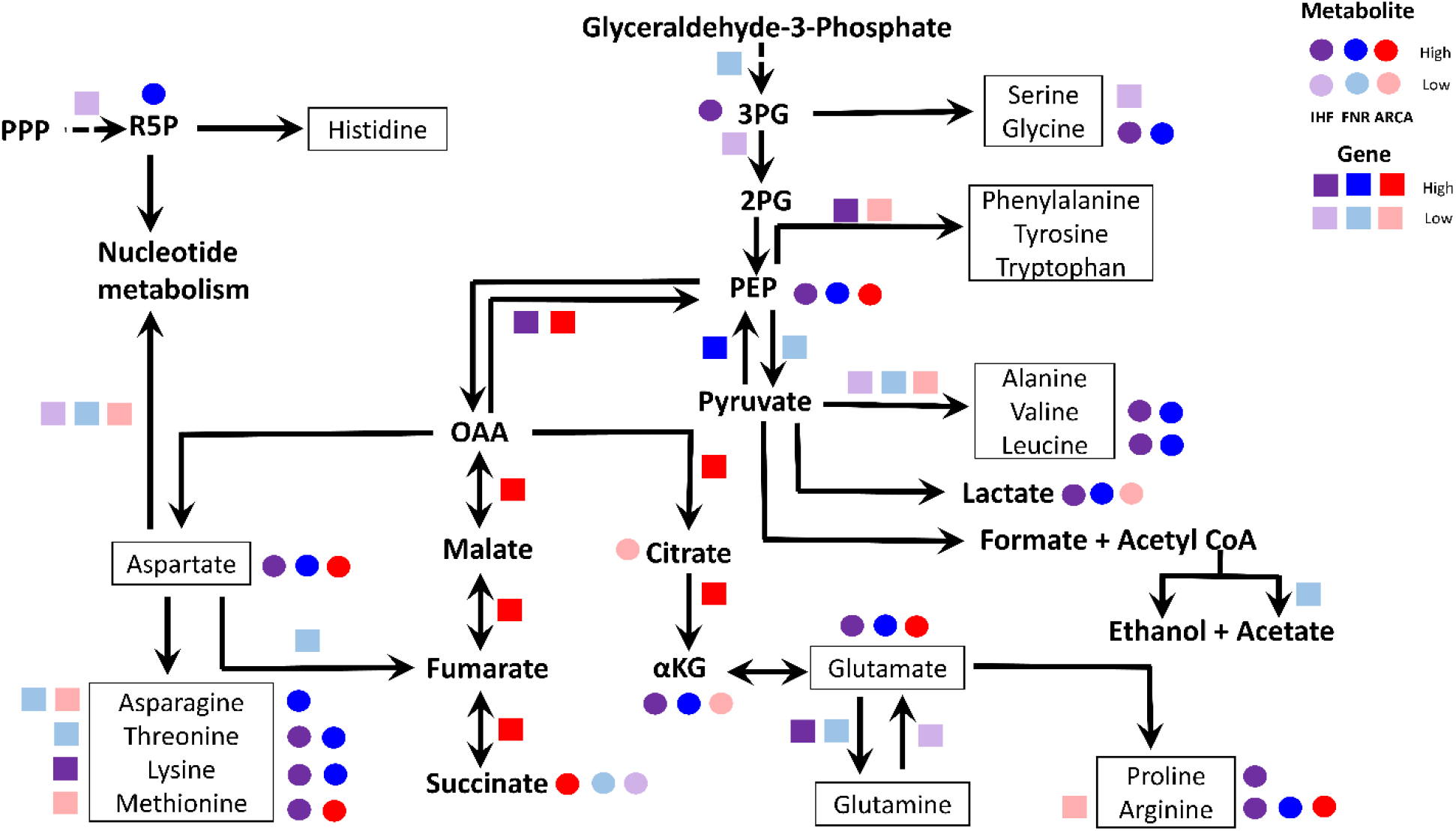
A schematic representation of the key steps in central carbon metabolic pathways affected by the deletion of either of the global regulators IHF, FNR and ArcA. The figure shows the unique pattern of regulatory control exerted on gene expression as well as metabolite levels. The genes are represented as coloured squares and metabolites are represented as coloured circles. Note: Succinate and Lactate represent exo-metabolite yields and not measured intracellularly. Darker colour represents upregulation whereas light colours represent downregulation respectively. Only statistically significant gene expression and metabolite levels are depicted. Abbreviations: 3PG — glycerate-3-phosphate; 2PG — glycerate-2-phosphate; PEP — phosphoenolpyruvate; αKG — α-ketoglutarate; PPP — pentose phosphate pathway; R5P — ribose-5-phosphate.

Next, we examined the TCA cycle intermediates such as α-ketoglutarate (αKG) and oxaloacetate (OAA) that are precursors for a large number of amino acids. It is well known that αKG that coordinates carbon and nitrogen balance by modulating glycolytic flux, accumulates during nitrogen limitation (33, 42). Indeed, both Δ*ihf* and Δ*fnr* mutant showed accumulation of αKG in contrast to Δ*arcA* that showed a reduction (Fig. 2 and Fig. S8). It is quite surprising that citrate and αKG levels are lower in Δ*arcA*, despite upregulation of *gltA* and *icd* genes, which might indicate reduced TCA cycle activity (15, 43).

The amino acids derived from αKG namely glutamate, proline and arginine, as well as the amino acids derived from OAA namely aspartate, lysine, methionine and threonine were found to be significantly higher in Δ*ihf* and Δ*fnr* mutant (Fig. 2 and Fig. S8, S9). In Δ*arcA*, concomitant with increase in glutamate and aspartate levels, we observed increased accumulation of arginine and methionine respectively that were in line with downregulation of methionine and arginine biosynthesis genes, reiterating feedback repression. Overall, such amino acid accumulations in all the mutants, could be, in part, due to the inability of the organism to efficiently utilize them towards protein biomass essential for optimal growth.

Noteworthy, our data indicates the downregulation of nucleotide biosynthesis genes (*nrdABEFG, purA*) in all the mutants, which succinctly explains the regulatory control on purine and pyrimidine metabolism (Fig. 2 and Fig. S7). Concomitant with increase in aspartate levels, we observed an increase in NAD^+^ levels in Δ*ihf* and Δ*fnr*, although *pntAB* gene involved in NAD^+^ dephosphorylation was downregulated solely in Δ*fnr* (Fig. S7). Additionally, we predict a higher redox state (NADH/NAD^+^ ratio) in Δ*arcA*, as inferred from lower level of NAD^+^ along with increase in gene expression of NAD^+^ phosphorylating *sthA* and decrease in non-proton translocating NADH dehydrogenase *ndh* (20, 43). To summarize, we observed direct and indirect effect on the metabolome due to the absence of global regulators, that corroborated with the upset in gene expressions in amino acid biosynthesis and other bottleneck reactions.

### Physiological characterization highlights the growth sub-optimality

Severe perturbation in metabolic processes due to regulator deletion, motivated us to examine their physiological effects on the organism. Characterization of the phenotype of the mutants revealed a reduction of ~16-25% (P < 0.05, Students t-test) in growth rate as well as glucose uptake rate compared to WT (Table S1). The decrease in growth rate was found to be positively correlated with decrease in glucose uptake rate (Pearson correlation coefficient (PCC) ~ 0.98, P < 10^-3^, Fig. S10A-C). Although glucose uptake rates were significantly lowered (P < 0.05, Student’s t-test), the *ptsG* gene that is solely responsible for glucose uptake was not differentially expressed in any of the mutants, indicating lack of transcriptional control on this gene by the regulators under the study. Nevertheless, the decrease in glucose uptake was in agreement with the accumulation of PEP observed across the mutants (PCC ~ −0.9, P < 0.05, S10A-C). We, further measured fermentation products or exo-metabolites namely formate, acetate, ethanol and lactate arising from pyruvate and succinate arising from PEP. We calculated the yields in each of the strains by normalizing their absolute secretion rates to their respective glucose uptake rates (Table S1). Succinate yield was found to vary among the mutants, wherein, Δ*fnr* showed ~55% decrease and Δ*arcA* showed ~33% increase, in comparison to WT (Fig. 2). Anaerobic succinate formation involves two steps wherein, 1) aspartate to fumarate formation is catalyzed by *aspA*, and 2) fumarate to succinate formation is catalyzed by *frd* genes coupled to NADH cycling *nuo* genes (30). Hence, changes in succinate yield can be attributed to the reduction in gene expression of *aspA* in case of Δ*fnr* and upregulation of *nuo* genes and reductive arm TCA cycle genes in Δ*arcA* (Fig. 1,2 and Fig. S9). Our findings of increased succinate yield *in ΔarcA* and reduced succinate yield in Δ*fnr* were consistent with previous studies (12, 16). Additionally, we obtained higher lactate yield in Δ*fnr*, that had strong correlations (PCC ~ 0.95, P < 0.005) with branched chain amino acids namely leucine and valine, both being derivatives of pyruvate (Fig. S10B). Next, we measured the ammonia uptake rate as an indicator of the nitrogen status of the organism. All the mutants showed a reduced ammonia uptake rate (P < 0.05, Student’s t-test) compared to the WT (Table S1) reflecting the nitrogen limitation faced by the organism, congruent with our gene expression and metabolite data (αKG, arginine, glutamate) as well. Thus, our data showed a profound effect on growth physiology, glucose and nitrogen import, thereby elaborating the system-wide effect of deletion of global regulators.

### Global transcriptional regulators control the translational and metabolic efficiency of the organism

Considering the key molecular features observed at the transcript and metabolite level that underscore the metabolic impairment due to the absence of each of the global regulators, we sought to quantitatively examine these changes at the proteome level that can directly account for the system-wide phenotypic effects such as growth rate and subsequent glucose uptake. Consequently, we recalled a proteome allocation model that quantitatively relates the cellular ribosome and metabolic proteome content to the growth of *E. coli* (4, 25–28). As explicitly outlined in their model (25), proteome sectors (Ø) are comprised of: R-sector defining the growth-rate dependent ribosomal protein sector, M-sector defining the metabolic protein sector and Q-sector defining the growth-rate independent core proteome sector (Fig. 3A). We defined an additional neutral proteome sector “U”, for the unused or unnecessary metabolic proteins, that help the organism hedge against altered environmental conditions but presumably poses a substantial proteomic burden under anaerobic fermentative conditions (Fig. 3A). These U-sector proteins defined here do not account for exogenously expressed proteins but rather endogenous proteins which can switch to being necessary subjected to nutritional or environmental conditions. However, these U-sector proteins reduce the proteome share available for R- and M-sector, effectively reducing the growth rate in analogy to inducible expression of exogenous proteins, devoid of any phenomenological parameters (25). Based on the proteome conservation law empirically outlined in the model (25), all the aforementioned sectors add up to unity:

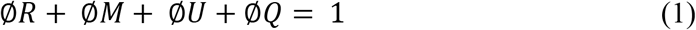

wherein the maximal growth-related total proteome fraction comprises of sum of R-, M- and U-sectors other than Ø*Q*,

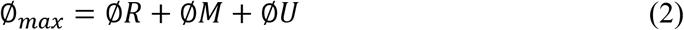

**Fig. 3.**
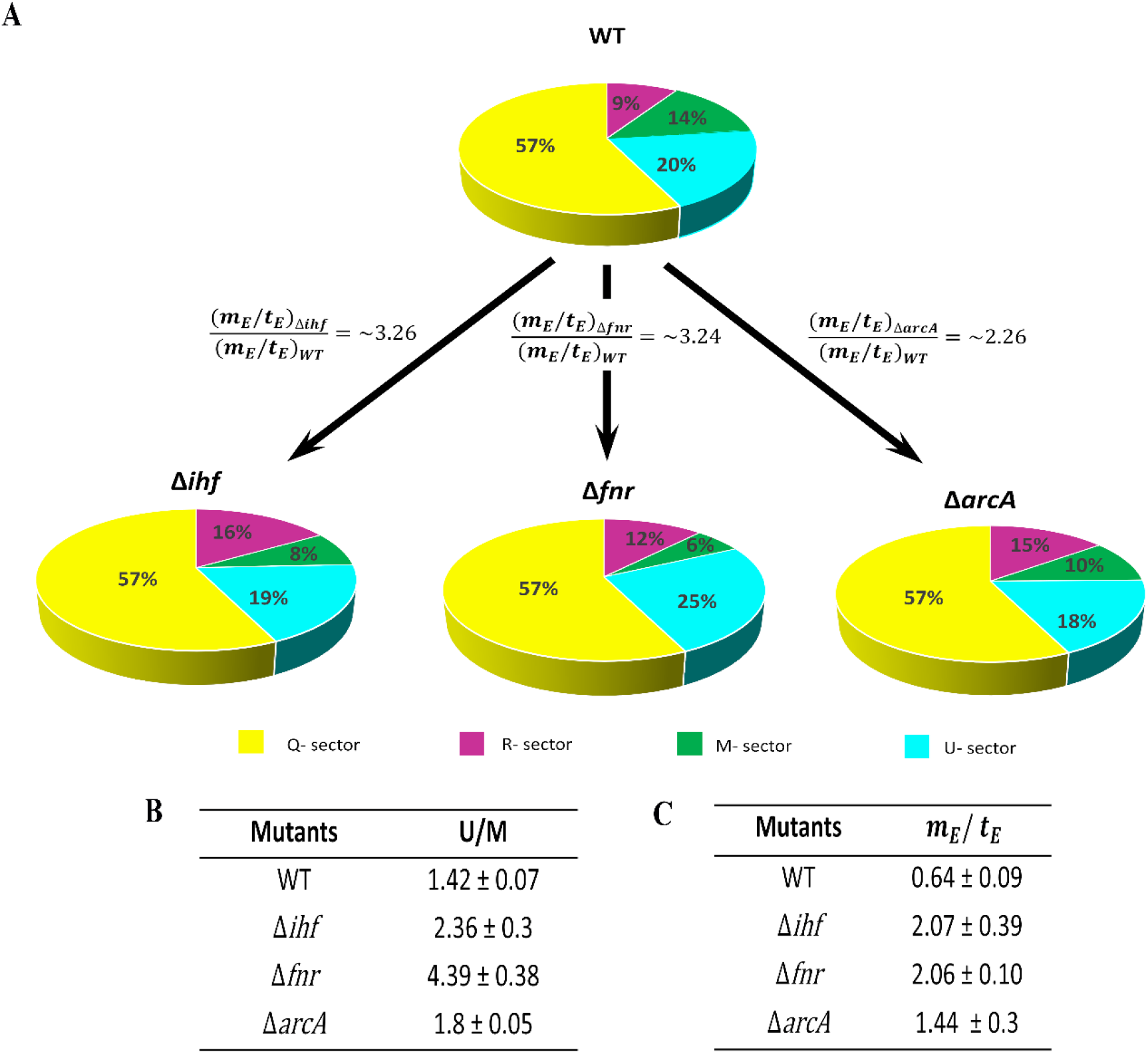
Proteome allocation in each of the strains. (A) A pie-chart depicting the quantitative distribution of the proteome sectors-R-sector, M-sector, Q-sector and U-sector as percentages, for the WT and the mutants. The figure indicates a high 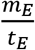 ratio represented as fold changes in each mutant compared to WT. (B) The U/M protein-coding transcriptome fraction ratio for each of the strains. (C) The 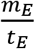 ratio derived for each of the strains.

Therefore,

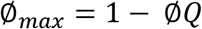

The R-sector was derived from the product of experimentally measured total RNA/total protein (R/P) ratio and empirically derived conversion factor ρ = 0.76 (25), in each of the strains (Fig. 3A). All the mutants in our study, showed a higher R/P value compared to the WT (Fig. S11) with Δ*ihf* showing the highest R/P. Assuming WT has an optimal R-sector, we consider that any excess above the optimal R-sector accounts for unused or unnecessary ribosomes in *E. coli*. We suggest that a higher R-sector might represent a compensatory countermeasure against the lowered growth of the mutants, as high ribosome supports faster growth (3, 4). This could be attributed to secondary growth effects mediated by metabolic regulator ppGpp (44), as majority of the ribosomal genes are devoid of binding sites for IHF, Fnr and ArcA. Notably, there exists a strong correlation of high ribosome content with lowered ppGpp levels (26). Interestingly, our analysis supports this notion in case of Δ*ihf* mutant, wherein DEGs that were reported to be positively regulated by ppGpp were downregulated (P < 10^-4^, Fisher exact test), possibly suggesting a lowered level of ppGpp in this mutant. This observation also reiterated the previously deduced overlap in ppGpp and IHF targets (24). However, such enrichments were found to be insignificant in case of Δ*fnr* and Δ*arcA* mutants. Studies have shown that despite any obvious enrichments observed for ppGpp, the R-sector could be modulated by simple supply driven circuits prevalent in exponentially growing cells (4, 45).

R-sector synthesis incurs a huge proteomic cost that reduces the available proteome resources towards M-sector. To evaluate the corresponding changes in M-sector, we sought to identify two key aspects, a) genes integral to glucose metabolism as a function of growth rate, and b) proteome fraction changes as deduced from commensurate changes in protein-coding transcriptome fractions. Towards this, we performed a simulation using ME (for Metabolism and macromolecular Expression) model that accounts for 80% of *E. coli* proteome (46, 47) (Supplementary Material and methods). From the simulation, we predicted the set of protein-coding genes that collectively accounted for M- and U-sector genes and subsequently mapped them to the combined list of DEGs from all the mutants. Further, we calculated the unnecessary/metabolic (U/M) ratio based on the protein-coding transcriptome fractions computed for each of the strains (Fig. 3B). As expected, we observed a consistent increase in the U/M ratio across the mutants compared to WT, with the highest for Δ*fnr* followed by Δ*ihf* and Δ*arcA*. Finally, using the Ø_*max*_ value (~ 43%) (26, 27), our Ø*R* and U/M ratio, we determined the Ø*M* as defined in Eq. (2).

For steady-state growth, the R-sector and M-sector proteome fraction have linear dependency on growth rate (26), described as:

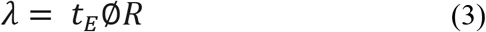

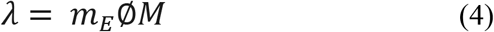

where, phenomenological parameter *t_E_* refers to translation efficiency (protein synthesis flux by R-sector proteome), phenomenological parameter *m_E_* refers to metabolic efficiency (metabolic flux attained by M-sector proteome) and *λ* refers to the growth rate (h^-1^) of the organism.

Further, by equating Eq. (3) and Eq. (4), we obtain the ratio of metabolic efficiency to the translation efficiency.

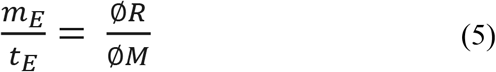

Hence, using experimentally derived growth rates and R-sector proteome fraction, and empirically derived M-sector proteome fraction, we computed the translational and metabolic efficiency in the mutants to quantify the extent of regulatory defect (Fig. 3C).

Intriguingly, deletion of global regulators resulted in an increase in 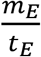 ratio, indicating a decrease in translation and increase in metabolic flux (Fig. 3A–3C). By normalizing the 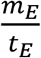 ratio for Δ*ihf*, Δ*fnr* and Δ*arcA* with WT ratio, we observed a concordant increase of ~3.26, ~3.24 and ~2.26 folds, respectively, that comprehensively quantifies the significance of these regulators in a WT strain. Collectively, the lowered *t_E_* and high *m_E_* as a result of imbalance in resource allocation, highlighted the inefficiencies of other growth-related physiological processes to counter the limitation in the absence of these global regulators (Fig. 3A).

### Physiological consequences of altered *m_E_* and *t_E_* on glucose uptake rates

Dynamic changes in growth pattern can be inferred from the abundances of the key intracellular metabolites (45, 48). Thus, the amino acids and precursor pool-sizes can be used as a proxy to describe the changes in *t_E_* and *m_E_* flux of the organism, given their linear negative association (Fig. S10A-S10C) with growth rate and glucose uptake rate in the mutants compared to WT. The accumulation of amino acids glutamate, aspartate and their derivatives observed consistently in the mutants owing to their inability to be incorporated into protein biomass, can be attributed to the reduction in *t_E_*. These accumulations along with changes in porin gene expression observed in the mutants, likely resemble an osmotic stress condition that agrees well with observed reduced translational capacity (49). Similarly, the increase in *m_E_* was indicated by accumulation of precursors namely PEP, αKG, OAA (speculated from aspartate levels) in the mutants.

Next, we probed how steady-state metabolite accumulations subjected to changes in growth rate, might affect glucose uptake rates in each of the mutants. In Δ*fnr* and Δ*ihf*, the intracellular concentration of αKG, when normalized to cell volume (2.3 μL/mg) (50), matches with its Ki (inhibition constant) value of 1.3 ± 0.1 mM for *ptsI* inhibition (42). *ptsI* codes for one of the components of the PTS system in *E. coli*, whose phosphorylation state determines the uptake rate of glucose. Thus, the higher intracellular concentration of αKG observed in Δ*fnr* and Δ*ihf*, can reduce the glucose uptake (PCC ~ −0.9, P < 0.05) by non-competitive inhibition on the PTS (42, 51). Similarly, OAA (speculated from aspartate levels), known to modestly inhibit the glucose uptake (51), was found to be present at higher concentrations in all the three mutants, with Δ*arcA* having the highest aspartate accumulation (PCC ~ −0.9, P < 0.05). Further, from the accumulation of leucine (PCC ~ −0.94, P < 0.01) and valine (PCC ~ −0.84, P < 0.05), increased lactate yield (PCC ~ −0.92, P < 0.01) and higher *m_E_* (Fig. 3C), we predict a high intracellular pyruvate pool in Δ*fnr*, that is known to adversely impact the glucose import (51). Moreover, it is known that the intracellular accumulated carbon-rich amino acids such as aspartate, glutamate and glycine via their degradation to α-keto acids can affect the glucose uptake rate (51). These observations, could in part, explain why Δ*arcA* showed slightly better glucose uptake compared to Δ*fnr* and Δ*ihf*. Additionally, extent of reduction in glucose uptake in each of the mutants corroborated with the reduction in M-sector and increase in U-sector (Fig. 3B) as well as increase in unused R-sector, given that glucose import and processing in minimal media condition involves significant metabolic proteome resources (7). Overall, each of the global regulators FNR, ArcA and IHF, show a distinct control over amino acid metabolism, key metabolic bottleneck steps and unnecessary gene expression, thereby optimizing the translational and metabolic machinery conducive for balanced growth.

## Discussion

Scrupulous proteome allocation in response to perturbations in the internal or external environment defines the growth physiology of *Escherichia coli* (1–4, 7). Anaerobic fermentation represents an energetically less favourable environmental condition characterized by high carbon uptake and slow growth (30). In this study, we addressed how global transcriptional regulators ensure efficient proteome allocation in a WT strain that enables it to attain an economical phenotypic outcome or foster robustness even in such unfavourable environmental condition. We explicitly demonstrated this using deletion effect of each of the global regulators FNR, ArcA & IHF on *E. coli* under anaerobic fermentation with glucose as a carbon source.

We elaborated the role of these regulators emphasizing their direct and indirect control over key metabolic processes fundamental for balanced physiological growth of *E. coli*. For instance, deletion of either FNR or ArcA or IHF ensued perturbation in key bottleneck steps of glycolysis, amino acid and nucleotide biosynthetic reactions, and derepression of aerobic TCA cycle and alternate carbon metabolic proteins. Using the growth-law theory (4, 25–28, 45, 49) and proteome-based ME model (38, 46, 47), we quantitatively accounted for this reduction in metabolic proteins and increase in unnecessary or hedging proteins, when an optimal WT strain is debilitated by global transcription regulator deletion. This reduction in metabolic proteome directly impacts the phenotype, thereby imposing a high translation program reflected as high ribosomal proteome to enable faster growth. However, owing to huge energetic cost associated with ribosomal synthesis itself, demand for increasing translation further constrains the proteome share for metabolic proteins. We illustrate that disruption of this inherent trade-off between metabolic and ribosomal proteomic investment ascribable to regulator deletion involved a lower translational and higher metabolic efficiency, thereby restraining growth (Fig. 4). Nonetheless, the lowered translational efficiency attempts to balance the perturbation in resource allocation by decreasing the synthesis of unnecessary and costly proteins although constraining the metabolic proteins as well. Concurrently, high metabolic efficiency, attempts to balance the perturbation in resource allocation by increasing the carbon import and processing activity in response to general limitations of carbon. The changes in these phenomenological parameters were manifested as accumulation of proteinogenic amino acids (glutamate, aspartate and their derivatives) and precursors (PEP, pyruvate, αKG & OAA), as evident from specific metabolic signatures in each of the regulator mutants. We suggest these amino acid accumulations to be a consequence of not being efficiently utilized for protein biomass, rather than accumulations that can occur due to protein degradation promoted by higher ppGpp levels evident in starvation conditions (52). Intuitively, fueling products (particularly α-ketoacids namely pyruvate, αKG & OAA) or building blocks (e.g. amino acids that on degradation generate α-ketoacids) have their own feedback mechanisms that can independently affect the translational and metabolic machinery (Fig. 4), limiting the potential of the system to attain growth rate similar to WT (40, 45, 48, 50, 51). Apart from carbon, we demonstrated that global regulators exhibit control over nitrogen homeostasis and sensing mechanism as evident from the induction of high-affinity nitrogen transporters or scavengers, metabolite profiles as well as the altered ammonia uptake. Overall, in the absence of global regulators, we propose that the balance or association between growth rate and glucose uptake rate is attenuated, making the organism incapable to attain fitter phenotype, unless forced upon a selection pressure (53, 54).

**Fig. 4.**
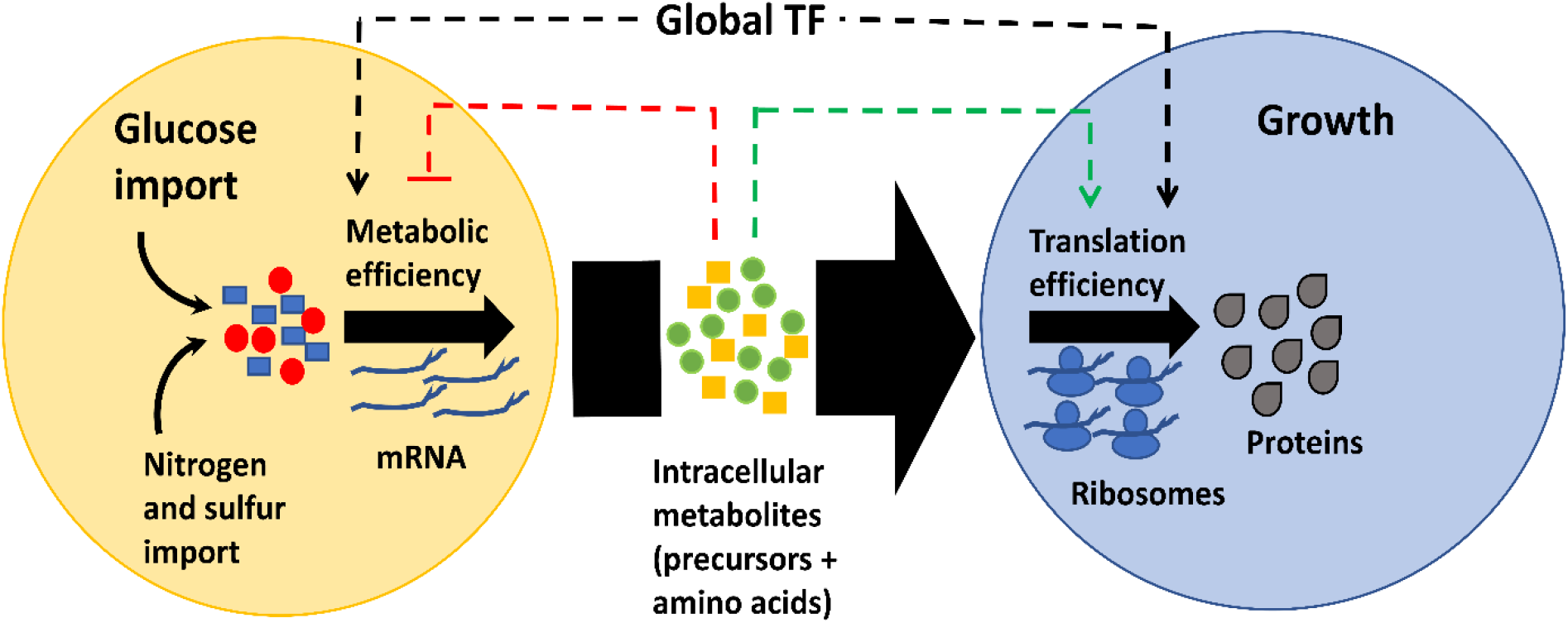
A model depicting global transcriptional factors (TF) control on the metabolic and translational machinery. Carbon and nitrogen sources, following their uptake are processed into intracellular precursors (yellow blocks) and amino acids (green circles) depending on the metabolic efficiency of the organism. These intracellular amino acids are then converted into proteins by ribosomes depending on the translation efficiency of the organism. The metabolic and translation efficiency of the organism is in turn feedback regulated by metabolites and genes under the purview of the regulators. Growth and glucose/carbon import have a strong linear association complemented by global regulators.

Under steady-state exponential growth conditions, protein levels are largely determined by their transcript concentrations albeit subjected to post-transcriptional and post-translational regulation (1), thus making it imperative to investigate the proteomic measurements together with metabolite interactions using mass spectrometry (55). However, traditional method of relative ribosome measurement has been reported to be in quantitative agreement with mass spectrometry or β-galactosidase assay data (28). Moreover, we anticipate that though variability can exist in gene expression studies, the metabolic and translational capacities simplified as fold changes would qualitatively be less variant to regulatory events at the mRNA or protein level. Nevertheless, our approach of relating the in-depth details to broad macroscopic properties addressed the general design principles concerning the existence and significance of global regulators in *E. coli*.

Mechanistically, this work elucidated the “complimentary” role of global transcriptional regulators with growth-rate dependent global machinery, predominant in exponentially growing cells. Using a well-established yet simplified theory (25), we were able to comprehensively demonstrate that absence of these global regulators ensued an upset in necessary and unnecessary proteome homeostasis thereby resulting in sub-optimal translational and metabolic machinery. Broadly, such analysis can be extended to other global regulators like CRP (27, 54), HNS (56), Lrp (57) and Fis (58), which tempts us to anticipate similar underlying mechanisms of regulation, presumably not confined only to glucose metabolism.

## Materials and Methods

Detailed materials and methods used for strain generation, physiological characterization, RNA extraction and mRNA enrichment, transcriptome data analysis, quantitative RT-PCR validation for RNA-seq, metabolomics, metabolomics data analysis, ME-model simulations, total RNA estimation and total protein estimation are provided as Supplementary Information.

### Data deposition

The RNA sequencing data and the processed files from this study are available at NCBI Geo (https://www.ncbi.nlm.nih.gov/geo/query/acc.cgi?acc) under accession no. GSE153906. The metabolomics data presented in this study is available at the NIH Common Fund’s National Metabolomics Data Repository (NMDR) website, the Metabolomics Workbench, https://www.metabolomicsworkbench.org where it has been assigned Project ID (PR000975). The data can be accessed directly via it’s Project DOI: (10.21228/M8NX1G).

## Acknowledgments

This work was supported by DST fellowship/grant(SB/S3/CE/080/2015) and DBT fellowship/grant(BT/PF13713/BBE/117/83/2015) awarded to K.V.V., and DST-WOSA grant (grant number: SR/WOS-A/ET-58/2017) awarded to S.S.

M.S.I. acknowledges Department of Biotechnology (DBT), Government of India for his fellowship (DBT/IIT-P/323). A.P. acknowledges Department of Science and Technology INSPIRE (DST-INSPIRE), Government of India for her fellowship (IF140914).

We would like to thank Dr. Aswin Sai Narain Seshasayee (NCBS, Bangalore) for his inputs in RNA-seq and constructive suggestions in the manuscript. We thank Edward Baidoo (Biochemist Research Scientist, Lawrence Berkley National Laboratory) for his valuable suggestions in metabolomics. We thank Colton Lloyd (Systems Biology Research Group, UCSD) for his inputs in ME-model simulations. We acknowledge Genotypic Technologies, Bangalore for library preparation and RNA sequencing. We thank Dr. Mayuri Gandhi (Research Scientist, SAIF, IIT Bombay) for providing the LCMS facility. We thank Prof. Pradeepkumar P. I. (IIT Bombay) for providing RT-PCR facility and Prof. Sarika Mehra (IIT Bombay) for providing vacuum concentrator for our use.

