## Supplementary information for "Global transcriptional regulators fine-tune the translational and metabolic machinery in *Escherichia coli* under anaerobic fermentation"

### Materials and Methods

#### Strain construction

We constructed *Δfnr, ΔarcA* and *Δihf* (*ihfA-ihfB*) knockouts in *E. coli* K12 MG1655 (CGSC#6300), by **λ**-Red mediated recombination using the plasmids pKD46, pKD13, pKD3 and pCP20 (1). The constructed strains were verified by PCR using the primers covering the region of interest as well as by Sanger sequencing. Glycerol stocks were made for each of the strains and stored at -80$^{o}$C.

#### Physiological Characterization

All omics characterizations were performed by growing cells in 400 mL M9 media (6 g/L anhydrous Na2HPO4, 3 g/L KH2PO4, 1g/L NH4Cl, 0.5 g/L NaCl + 2 mM MgSO4 + 0.1 mM CaCl2) (25) with 2g/L glucose in a 500 mL capacity bioreactor (Applikon). Briefly, cells from glycerol stocks were plated out on LB + Kan (Kanamycin 50 µg/mL) agar plate and a single colony was inoculated in LB media. A fixed volume of 100 µL cells were used to inoculate 50 mL preculture M9 media with 4 g/L glucose which was grown overnight in shake flask at 250 rpm in a 37$^{o}$C (Eppendorf) incubator. The preculture cells, still in exponential phase, were centrifuged and washed with M9 (no carbon source), and inoculated in a bioreactor containing 400 mL M9 media with 2g/L glucose such that the start OD of all the cultures were ~ 0.07 OD. The temperature of the bioreactor was maintained at 37$^{o}$C, the stirrer speed was 150 rpm and the pH of the media was maintained at pH 7.2 using 1M NaOH. The pH was continuously monitored using a pH probe.

Dissolved oxygen (DO) levels were monitoring continuously using polarographic dissolved oxygen probe and maintained at zero by constantly sparging nitrogen. The growth rate was measured from the slope of the linear regression line fit to natural logarithm of the optical density (O.D.) values at 600 nm wavelength versus time plot, ln(OD@600) vs. time (in hours), during the exponential growth phase. The dry cell weight (DCW) was experimentally determined for each strain across the exponential phase such that 1.0 O.D. @600nm corresponds to 0.44 gDCW*$h^{-1}$. Three biological replicates (n=3) were considered for all phenotypic characterizations. To determine the rate of glucose uptake as well the rate of secretion of mixed acid fermentation metabolites (acetate, lactate, pyruvate, succinate, formate, ethanol), samples were collected throughout the exponential phase (2) which were then centrifuged. The supernatants were used to determine the concentrations using HPLC (Agilent 1200 Series) equipped with Bio-Rad Aminex HPX-87H ion exclusion column, with 5mM $H_2SO_4$ as the mobile phase. The column temperature was maintained at 50$^o$C and the flow rate at 0.6 mL/min. The supernatants were also used to determine the ammonia uptake rate using an enzyme-based assay (Sigma #AA0100). Given that pyruvate, succinate and lactate are also secreted by the bacterial culture in substantial amounts, these metabolites were confined to exo-cellular measurements. Yields of biomass and mixed acids fermentation products were calculated by normalizing their rates with glucose uptake rates (g/g Glucose).

**RNA extraction and enrichment of mRNA**

The RNA extraction from two biological replicates of each strain were performed (n = 2). Briefly, the cells were grown anaerobically till mid- exponential phase after which 50 mL

(0.35-0.4 OD) cells were harvested by centrifugation. The TRIzol-chloroform method was used to extract to total RNA (3, 4) as detailed below. DNase treatment was done to remove any genomic DNA contamination. After DNase treatment, an initial enrichment of total RNA using MegaClear Kit (Ambion) was performed as per manufacturer's instruction. Enrichment of mRNA was done using the MicrobExpress Kit following the manufacturer's protocol and the quality and integrity was checked using BioAnalyzer. Paired-end, strand specific libraries for RNA sequencing were prepared using NEBNext Ultra Directional RNA library kit and the sequencing was carried out on HiSeq 4000 Rapid Run Mode using 2x150 bp format, at Genotypic Technologies, Bangalore.

**Transcriptome data analysis**

Raw reads filtering, trimming, mapping and alignment were performed as reported previously (5) taking into consideration the strand specific paired end reads. The error tolerance for filtering low quality reads and adapter sequences was set to 20% and trimmed reads with <50 bases were excluded. The annotations for 4466 genes excluding the rRNA, tRNA and sRNA genes were extracted from Ecocyc database (version 21.5) (6). Raw counts from EdgeR (7) were then used for analysis of differential gene expression (DGE), after filtering based on genes with reads $< 0.5$ counts per million (cpm). Gene showing $\geq$ 2-fold change in expression and adjusted $P < 0.05$ (Benjamini-Hochberg) were used for all further analysis.

Enrichment of up-regulated and down-regulated DEGs were done separately for regulators FNR, ArcA and IHF using information available in Ecocyc (6) and statistical significance was performed using Fisher Exact test. Only enrichments with $P < 0.01$ were considered for further analysis.

For sigma factor enrichment analysis, the up-regulated and down-regulated DEGs were separately enriched based on sigma factor targets using data available in EcoCyc and Regulon DB (10). The upregulated and downregulated genes under each regulator were then validated using hypergeometric test in R (P-value < 0.05). Only those sigma factors which regulated atleast 5 genes from either the upregulated or downregulated DEGs in each of the mutants, were retained for this over-representation analysis.

The up-regulated and down-regulated DEGs were separately enriched for metabolic pathways using KEGG pathway classification (8) as defined in Proteomaps (www.proteomaps.net) and the mapped genes were represented as Voronoi treemaps (version 2.0) (9). A hypergeometric test with P-value correction using Benjamini Hochberg, was applied to determine the significance of the upregulated and downregulated genes within each pathway in R (R Core Team 2019) (26). For each upregulated and downregulated pathway, we arbitrarily choose atleast ten DEGs to be considered for significance analysis. DEGs not having any assigned “Accession ID” (Ecocyc Version 21.5) such as phantom genes were excluded from the above analysis.

The ppGpp enrichment analysis was performed on up-regulated and down-regulated DEGs separately using data available in EcoCyc and statistical significance assessed using Fisher’s Exact Test. Only enrichments with $P < 0.01$ were considered for further analysis.

**Quantitative RT-PCR validation for RNA-Seq**

RT-PCR was performed with the Agilent AriaMx machine using the PowerUp SYBR Green PCR Master Mix to validate the RNA-Seq data. *rpoB* was used as internal control to normalize the RTqPCR data. DNAse treated Total RNA to cDNA conversion was performed using Superscript III Reverse Transcriptase following the manufacturer's instructions. All experiments were performed in biological duplicates and technical triplicates (n=6). $2^{-\Delta\Delta Ct}$ method described previously (11) was used to quantify expression fold changes. We observed a strong correlation (> 0.93) between qRT-PCR and RNA Seq data.

**Metabolomics**

Metabolite samples were harvested from anaerobically growing cells in mid-exponential phase. Three biological and two technical replicates (n=6) were harvested as reported previously (5) with minor changes specific to anaerobic conditions. A Fast-Cooling method was used to quench the harvested cells as reported previously (12, 13). Briefly, ~15 mL culture (> ~5 O.D. cells) was rapidly poured into 5 mL chilled M9 (without glucose) in a pre-cooled 50 mL falcon tube. To rapidly bring the temperature of the sample tube down to $0^{o}C$, the tube was dipped in liquid nitrogen for 10 secs with vigorous agitation with the help of a digital thermometer, to prevent ice crystal formation. Samples were then immediately centrifuged at $0^{o}C$, 7800 rpm for 7 min. The supernatant was discarded and the pellet was snap frozen in liquid nitrogen and stored at -80$^{o}$C until metabolite extraction was done.

For metabolite extraction, 7:3:5 methanol: chloroform: 2% ammonium hydroxide was used as described previously (5, 13). $^{13}$C labelled *E. Coli* extracts were used as internal standards that were generated separately in aerobic flask conditions using the wild-type strain. The labelled extracts were used in the quantification of key metabolite pool sizes using isotope-based dilution method (14). All the extracted samples were spiked with a fixed volume of internal standard taken from the same batch at an early stage of extraction. The volume of the pooled internal standard added to the samples was accepted only if the external $^{12}$C peak height (concentration similar to samples) and internal $^{13}$C standard peak height differed less than 5-fold (15). The LC-MS/MS settings (5) and chromatographic conditions (5, 16) were maintained as reported previously. Cleaning and maintenance of LCMS was performed (17) before the actual setup. The ESI was operated in positive [M+H]+ and negative [M-H]- mode separately. MS1 parent ion was used for quantification purpose. The MS2 setting was used for secondary validation of metabolites including the $^{12}$C chemical standards.

**Metabolomics data analysis**

The raw files generated from the machine were processed using the software package Xcalibur 4.3 (Thermo Fisher Scientific) Quan Browser as reported in our previous study (5). Quantitative analysis was performed wherein the peak heights of precursor ions with signal/noise (S/N) ratio more than 3 and less than 5 ppm error were considered. Metabolite height ratio was obtained after normalizing peak heights of the samples to the peak height of the internal standards. To identify the concentrations, serial dilutions of $^{12}C$ chemical standards (mix of 40 metabolites) supplemented with the fixed volume (as in samples) from the same batch of $^{13}C$ labelled internal standards, were used to generate a calibration curve in the range of 0.781 µM to 50 µM. These standards were run in biological duplicates (n=2) for the above-mentioned concentration range in the positive and negative mode separately. All metabolite concentrations were within the calibration curve range and those which do not fall in this range were individually checked for that particular concentration to assess whether they lie within the limit of quantification (LOQ) (S/N = 10).

MetaboAnalyst (18) was used for identifying statistically significant metabolites. Biomass normalized concentrations on metabolites were g-log transformed prior to analysis. Missing value imputation were performed using SVD impute function in MetabolAnalyst. The absolute concentration of metabolites is expressed as µmol/gDCW. Only those metabolites with false discovery rate (FDR) value $< 0.05$ (Students t-test) were considered for further analysis.

We sought to identify the specific pattern of precursor or amino-acid correlations with the glucose uptake and growth rate across the WT and mutant condition, given the physiological state of the system perturbed due to a regulator deletion. Towards this, we performed pairwise Pearson correlation analysis with statistical significance ($P < 0.05$) between metabolites and phenomic features such as glucose uptake, growth rate etc. in each of the mutants compared to WT in R. For this analysis, only features/metabolites found to be statistically significant (FDR $< 0.05$ from Students t-test) in each mutant compared to WT were considered for correlation analysis. The concentration (n=6) of technical and biological replicates of metabolites were clubbed (n=3) to make it comparable with phenomic features (n=3). These concentrations or rates were log2 normalized before assessing the correlation in R (R core 2019) (26).

**ME- model (Metabolism and macromolecular Expression) simulations**

We assessed the utilized ME and non-utilized ME protein coding fractions as reported in our previous study (5), using an *E. coli* ME-model (19, 20). The simulation involved constraining the glucose uptake rates in the range from zero to unbounded glucose (5, 21, 22), with an additional constraint on oxygen that was set to zero, with the maximization of growth rate as objective function. Genes predicted to have a reasonable protein translation flux in any of the simulations were classified as "utilized ME"; genes within the scope of the ME-Model that showed no expression or very low expression (protein translation flux $< 10^{-15}$) were classified as "non-utilized ME". Two assumptions were considered in this analysis - 1) the utilized ME and non-utilized ME correspond to metabolic "M" sector and unnecessary/unused "U" sector genes respectively and 2) the increase in these protein coding transcriptome fraction correspond to increase in proteome fractions. DEGs utilized in the simulation correspond to genes that were differentially expressed (adj-P value < 0.05, aFC >=2) in atleast one condition (*Δfnr* vs WT, *ΔarcA* vs WT, *Δihf* vs WT). We mapped the DEGs in each of the mutants compared to WT to the ME model predicted protein-coding genes to collectively account for M-sector and U-sector genes. This mapping was done utilizing the combined list of DEGs from all the mutants, to make it comparable across the WT and all mutant strains. Additionally, DEGs outside the scope of ME model were further enriched using KEGG pathway classification and added to M-sector and U-sector gene list based on manual annotation using Ecocyc (6,8, 9). Raw counts (genes < 0.5 cpm reads were not considered) for all the 4466 genes in WT and the mutants were used for calculation of transcript per million (TPM) (23). TPMs were assigned to all the M-sector and U-sector genes annotated to DEGs. The transcriptome fraction was calculated using the product of TPM and gene length divided by the sum-product of these calculated over the sector specific genes. Finally, we summed the transcriptome fractions of all M-sector and U-sector genes independently (22). The U/M ratio was calculated by dividing these protein-coding trancriptome fractions of U-sector and M-sector.

**Total RNA estimation**

Total RNA extracted from the cells using TRIzol-chloroform method as per the manufacturer's instructions. Briefly, 50 mL cells were harvested from the bioreactor around 0.35-0.4 O.D. Cells were pelleted by centrifugation at 4 $^{o}$C. The pellets were stored at -20$^{o}$C until utilized for further processing steps. The pellet was snap frozen in liquid nitrogen followed by addition of 300 µL TRIzol. The pellet was homogenized using hand-held pestle for not more than 2 mins. To this homogenized pellet, 700 µL TRIzol was added, vortexed and kept on ice for 15 mins. Further, extraction was done using chloroform (300 µL) and centrifuged to separate the protein, DNA and RNA layers. The RNA layer was then precipitated using isopropanol as per the instructions provided in TRIzol method. Next, RNA was centrifuged at 4 $^{o}$C and the RNA pellet was washed with 70% ethanol at room temperature, making sure there are no traces of ethanol. The pellet was dried by inverting the tubes and to the dried pellet 30-40 µL DNAse-RNAse free water was added. This reconstituted sample was incubated in a dry bath at 55 $^{o}$C for 10 mins, brought to room temperature and then stored as aliquots at -80 $^{o}$C until use. The total RNA was checked for its purity and integrity by running on an agarose-formaldehyde gel and Bioanalyzer and its concentration was estimated using Nanodrop.

**Total Protein estimation:** The total protein quantification using 1.8 mL culture around 0.35-0.4 OD, was based on the Biuret method described previously (24).

### Supplementary Figures

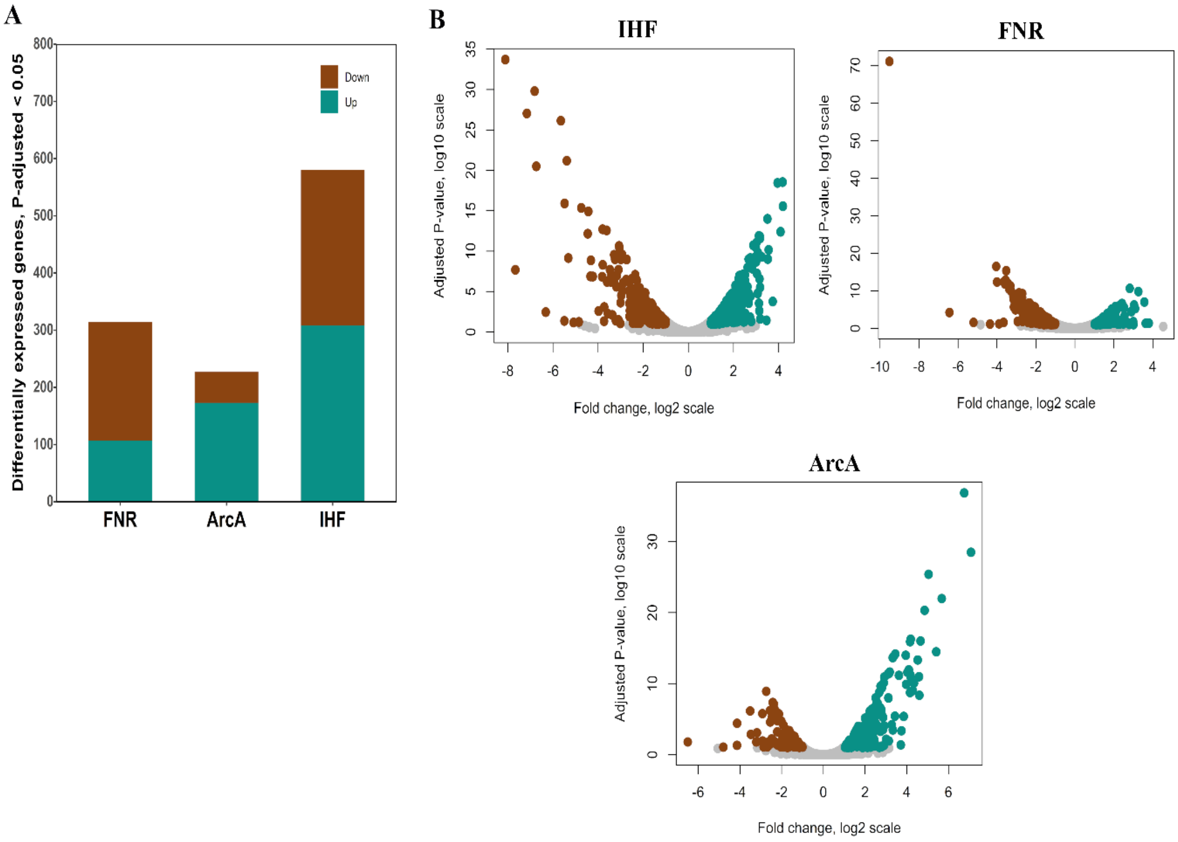


**Fig. S1.** Transcriptome comparison of the regulator mutants compared to WT. (A) Stacked plots showing the number of DEGs in all the mutant strains. (B) Volcano plot of the DE genes for Δ*ihf*, Δ*fnr* and Δ*arcA* compared to WT, depicted as adjusted P (adj-P) value (log10 scale) vs Fold change (log2 scale). The downregulated genes are shown in brown and the upregulated genes are shown in cyan.

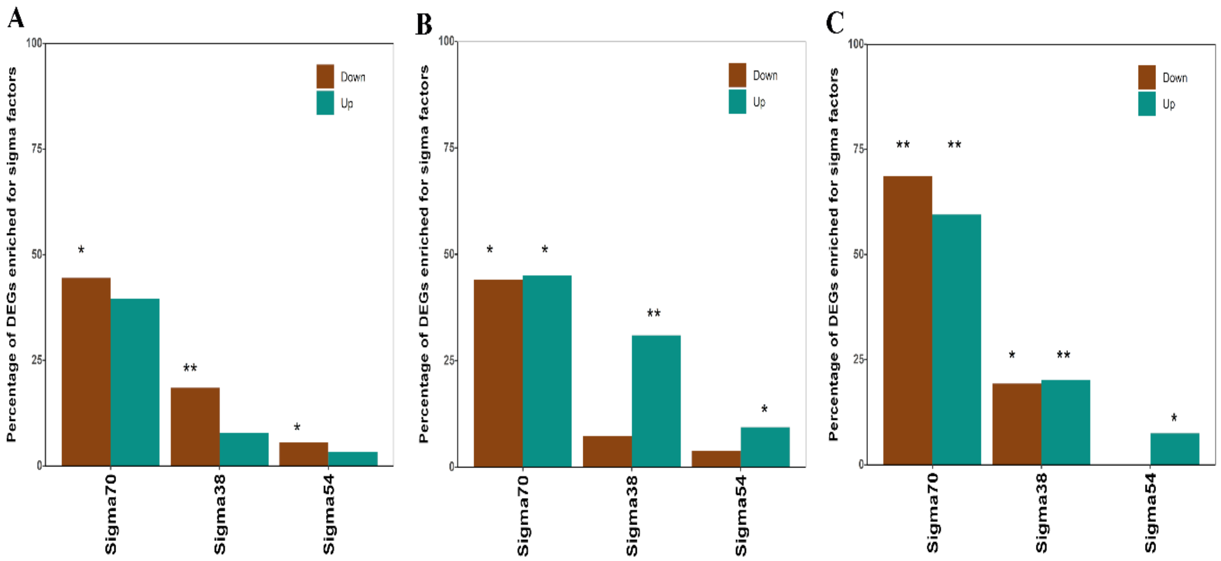


**Fig. S2.** Enrichment of DEGs under regulation of sigma factors in each mutant compared to WT; (A) Sigma factor enrichment in Δ*ihf*. (B) Sigma factor enrichment in Δ*fnr*. (C) Sigma factor enrichment in Δ*arcA*. The brown bars and the cyan bars indicate the fraction of downregulated and upregulated genes, respectively. Significant increase or decrease is denoted by asterisks: one asterisk indicates $P < 0.05$ and two asterisks indicate $P < 10^{-4}$.

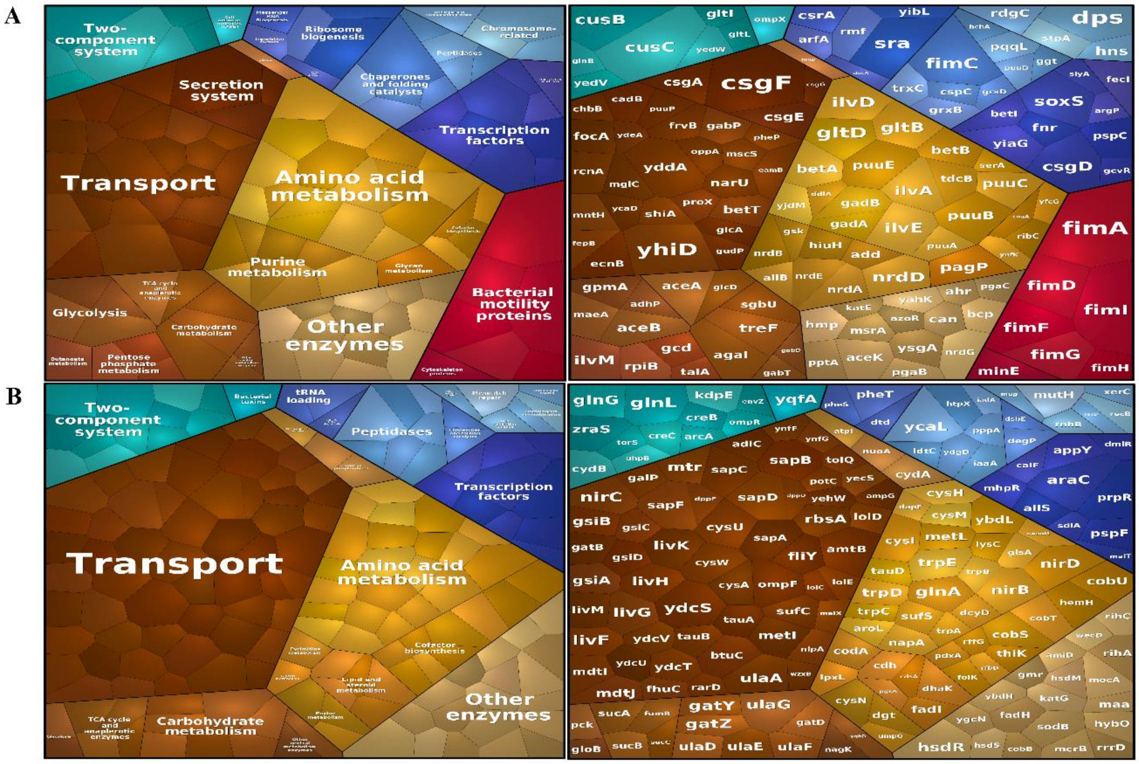


**Fig. S3.** The Voronoi maps showing the metabolic pathways and the genes within each pathway, enriched by KEGG classification in Δ*ihf* compared to WT. (A) The downregulated metabolic pathways and the genes within each pathway. None of the pathways were found to be significantly down-regulated. (B) The upregulated metabolic pathways and the genes within each pathway. Amino acid metabolism (adj-P < 0.05) and Transport (adj-P < $10^{-3}$) were found to be significantly upregulated. The size of the hexagon within each category is directly proportional to the absolute fold change observed for the genes. The colour of the hexagon denotes the specific pathways classified by KEGG.

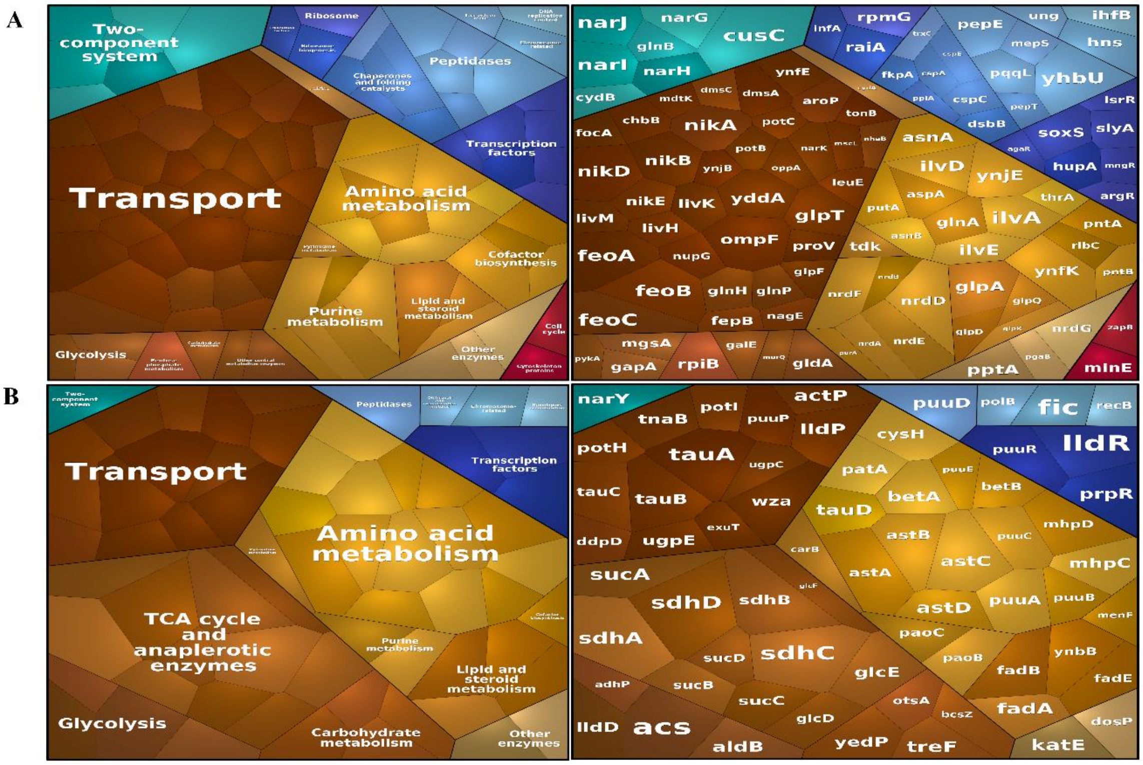


**Fig. S4.** The Voronoi maps showing the metabolic pathways and the genes within each pathway, enriched by KEGG classification in Δ*fnr* compared to WT. (A) The downregulated metabolic pathways and the genes within each pathway. Transport (adj-P < 0.002) was found to be significantly downregulated. (B) The upregulated metabolic pathways and the genes within each pathway. Amino acid metabolism (adj-P < $10^{-3}$), and TCA cycle (adj-P < $10^{-8}$) were found to be significantly upregulated. The size of the hexagon within each category is directly proportional to the absolute fold change observed for the genes. The colour of the hexagon denotes the specific pathways classified by KEGG.

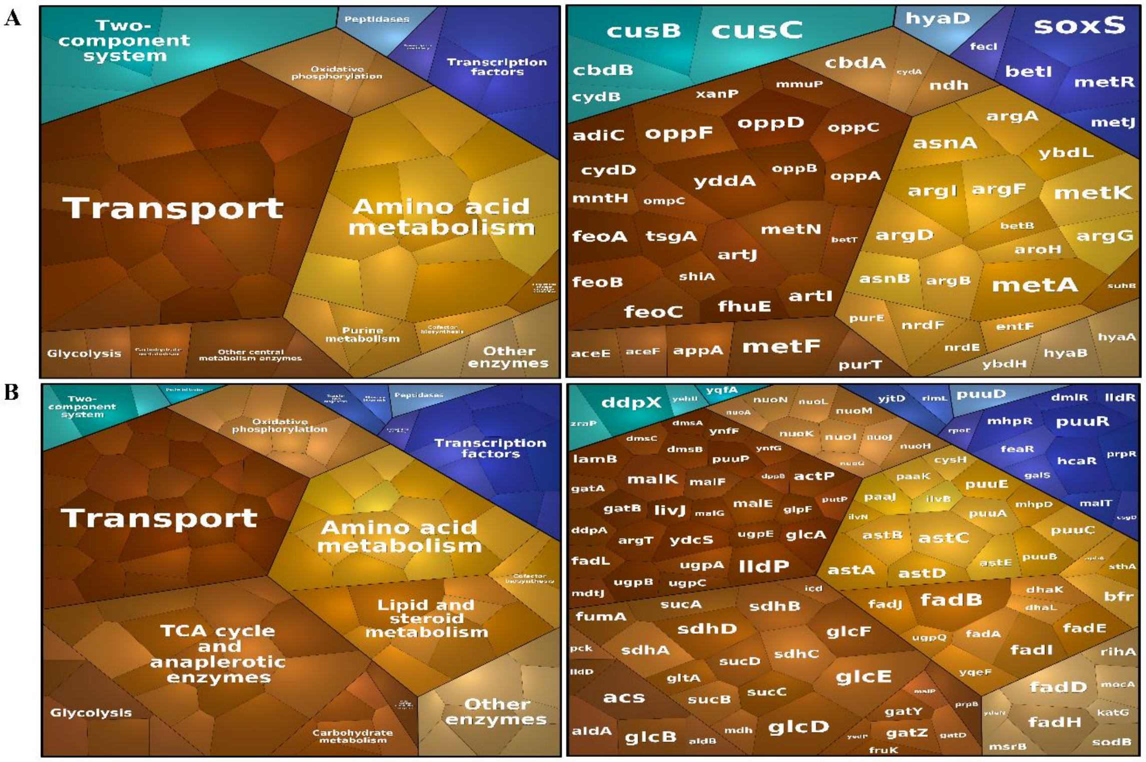


**Fig. S5.** The Voronoi maps showing the metabolic pathways and the genes within each pathway, enriched by KEGG classification in Δ*arcA* compared to WT. (A) The downregulated metabolic pathways and the genes within each pathway. Amino acid metabolism (adj-P < $10^{-3}$) and Transport (adj-P < $10^{-3}$) were found to be significantly downregulated. (B) The upregulated metabolic pathways and the genes within each pathway. Amino acid metabolism (adj-P < 0.0152), TCA cycle (adj-P < $10^{-10}$), Transport (adj-P < 0.012), Oxidative phosphorylation (adj-P < $10^{-5}$), and Lipid metabolism (adj-P < $10^{-3}$) were found to be significantly upregulated. The size of the hexagon within each category is directly proportional to the absolute fold change observed for the genes. The colour of the hexagon denotes the specific pathways classified by KEGG.

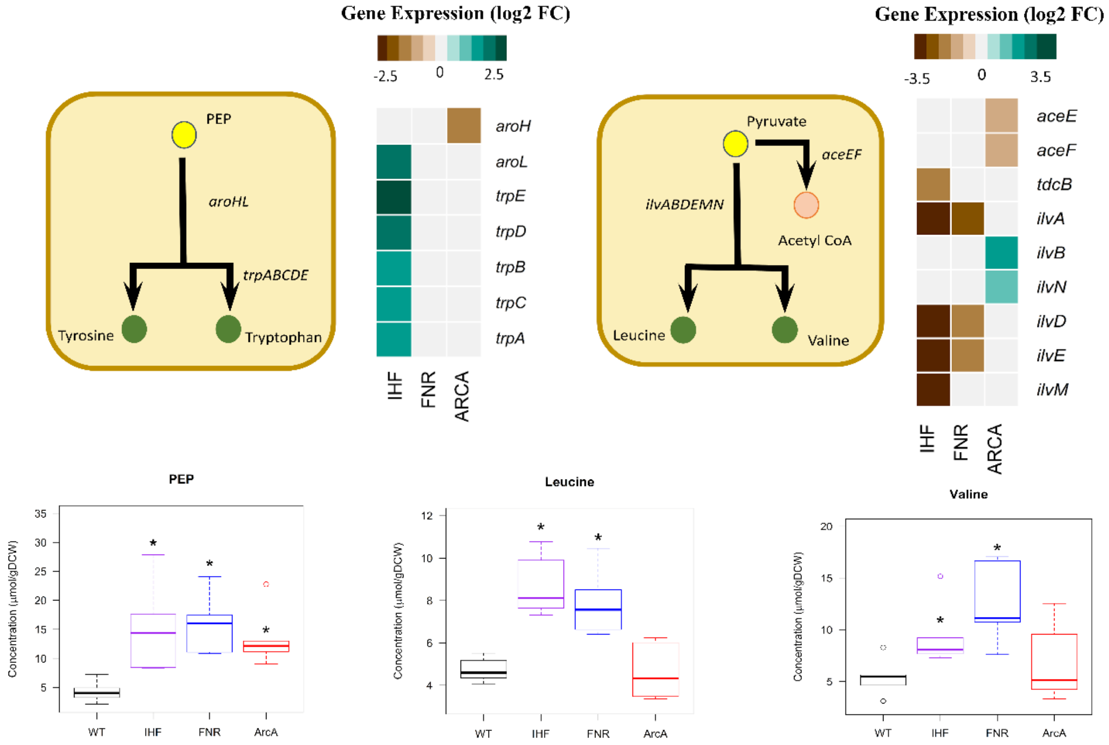


**Fig. S6.** Integrated transcriptomics and metabolomics analysis at PEP and pyruvate node. The ball and stick representation of the pathway annotated with genes is obtained from Ecocyc, wherein yellow color represents the precursors PEP and pyruvate and green color represents amino acids. Gene expression profile of DEGs altered in the pathways depicted as heatmaps, are obtained by comparing each of the regulator mutants (Δ*ihf*, Δ*fnr*, Δ*arcA*) with WT. Expression values are obtained from average of two biological replicates (n=2) expressed as log2 Fold change. Metabolite concentrations are obtained from average of three biological and two technical replicates (n=6) expressed as µmol/gDCW.

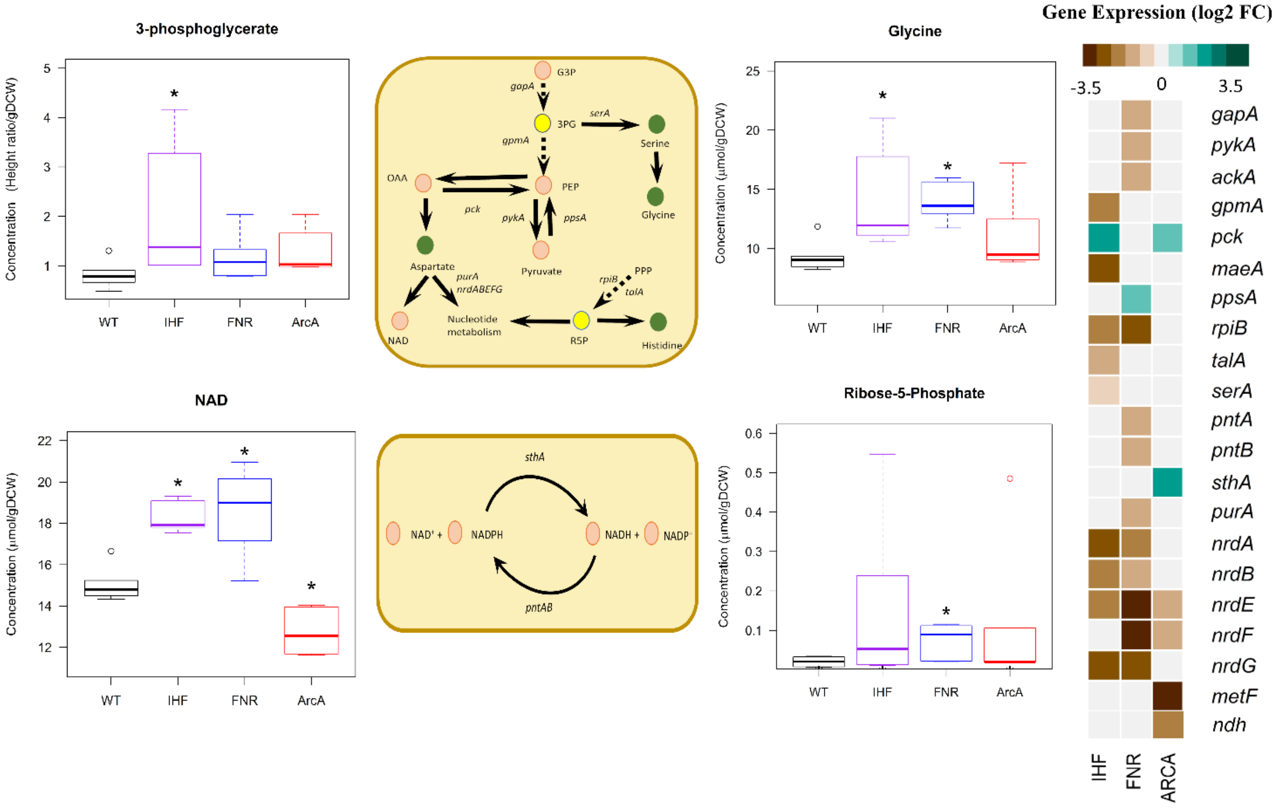


**Fig. S7.** Integrated transcriptomics and metabolomics analysis at 3PG and R5P node. The ball and stick representation of the pathway annotated with genes is obtained from Ecocyc, wherein yellow color represents the precursor 3PG and R5P and green color represents amino acids. Expression profile of DEGs altered in the pathways depicted as heatmaps, are obtained by comparing each of the regulator mutants (Δ*ihf*, Δ*fnr*, Δ*arcA*) with WT. Gene expression values are obtained from average of two biological replicates (n=2) expressed as log2 Fold change. Metabolite concentrations are obtained from average of three biological and two technical replicates (n=6) expressed as µmol/gDCW.

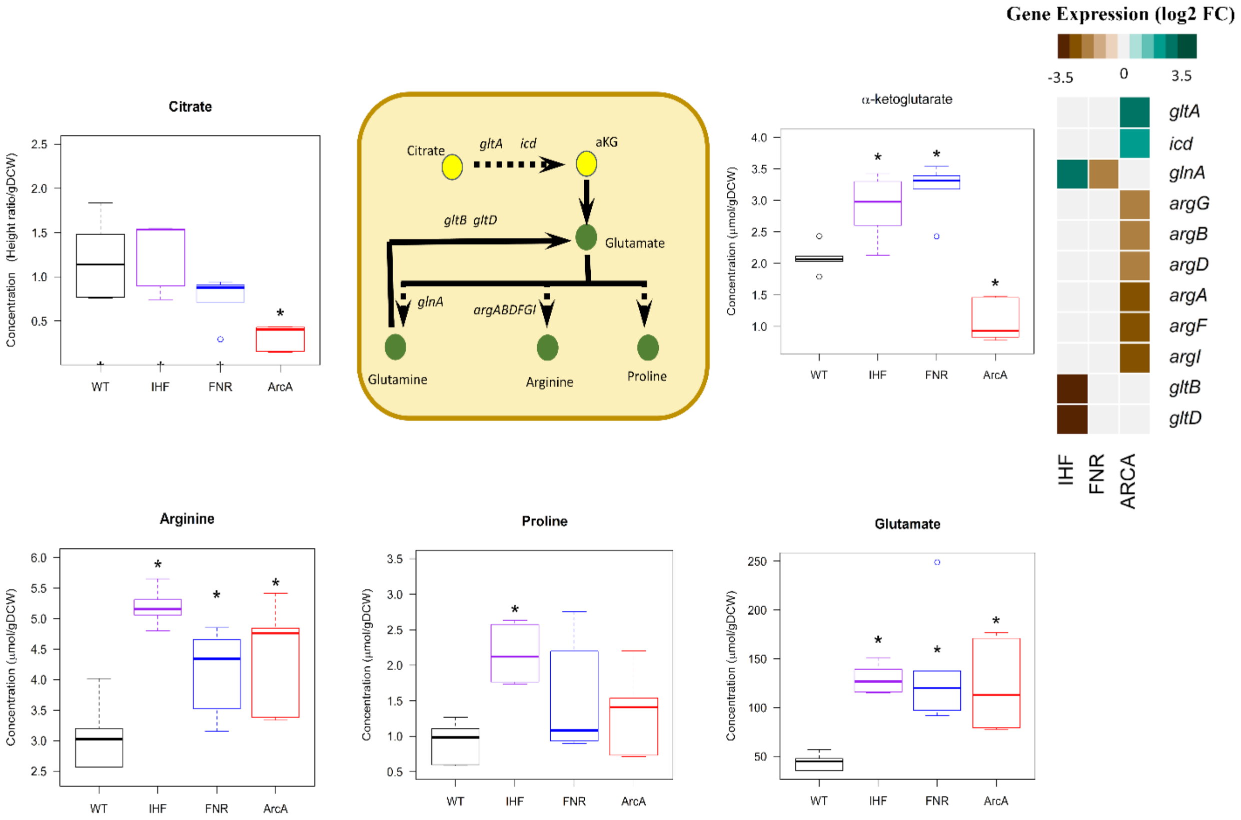


**Fig. S8.** Integrated transcriptomics and metabolomics analysis at citrate and αKG node. The ball and stick representation of the pathway annotated with genes is obtained from Ecocyc, wherein yellow color represents the precursors citrate and αKG and green color represents amino acids. Expression profile of DEGs altered in the pathways depicted as heatmaps, are obtained by comparing each of the regulator mutants (Δ*ihf*, Δ*fnr*, Δ*arcA*) with WT. Gene expression values are obtained from average of two biological replicates (n=2) expressed as log2 Fold change. Metabolite concentrations are obtained from average of three biological and two technical replicates (n=6) expressed as μmol/gDCW.

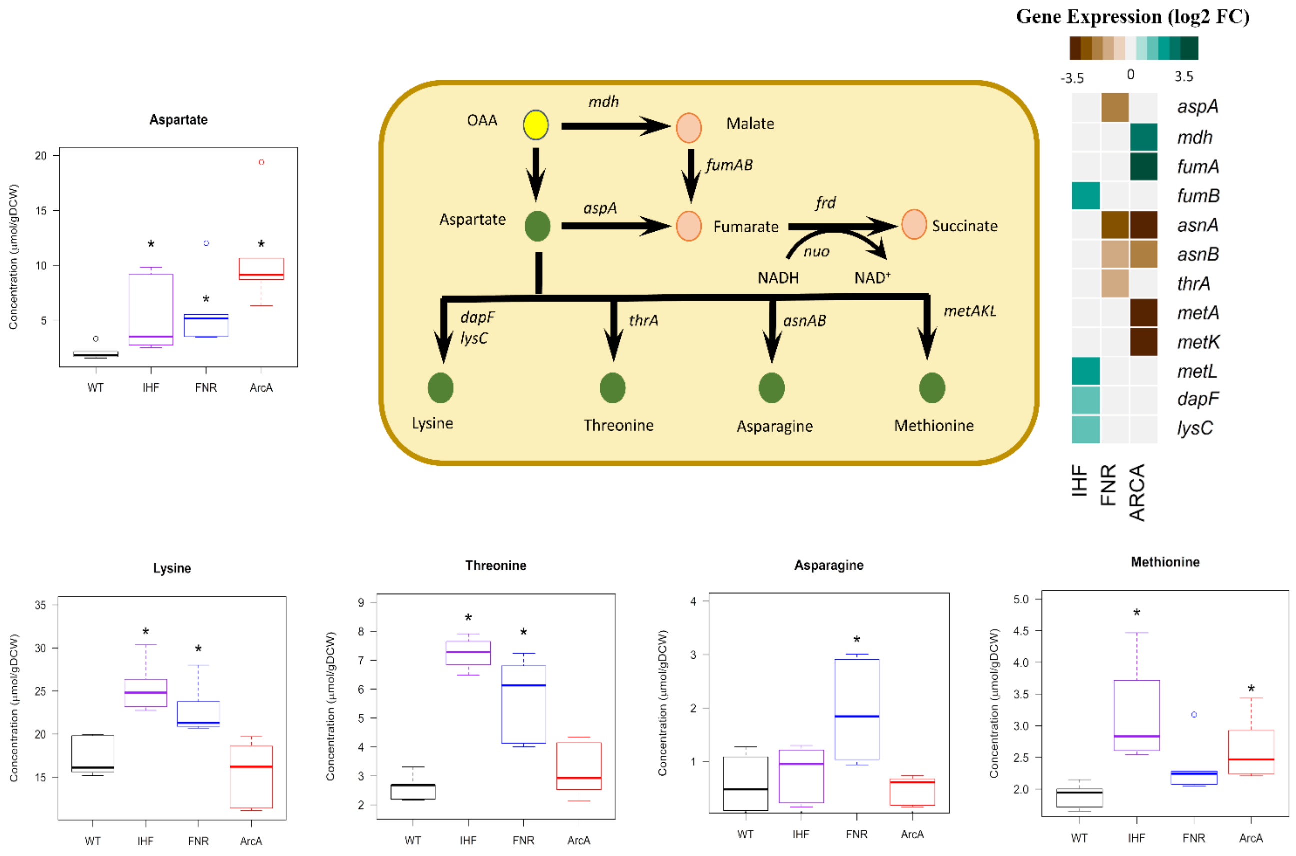


**Fig. S9.** Integrated transcriptomics and metabolomics analysis at OAA node. The ball and stick representation of the pathway annotated with genes is obtained from Ecocyc, wherein yellow color represents the precursor OAA and green color represents amino acids. Gene expression profile of DEGs altered in the pathways depicted as heatmaps, are obtained by comparing each of the regulator mutants (Δ*ihf*, Δ*fnr*, Δ*arcA*) with WT. Expression values are obtained from average of two biological replicates (n=2) expressed as log2 Fold change. Metabolite concentrations are obtained from average of three biological and two technical replicates (n=6) expressed as µmol/gDCW.

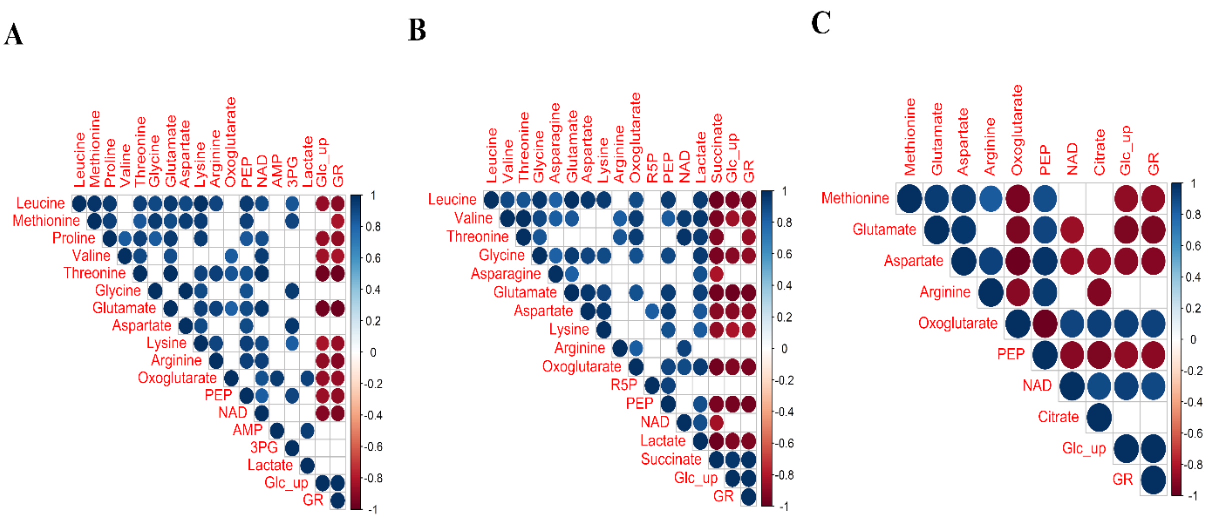


**Fig. S10.** Pairwise Pearson correlation plot of log2 normalized metabolite concentrations as well as glucose uptake rate (Glc_up) and growth rate (GR) for A) Δ*ihf* with WT, B) *Δfnr* with WT and C) Δ*arcA* with WT. Only the significantly altered metabolites in each mutant compared to WT were considered for the analysis. Succinate and lactate represent log2 normalized exo-metabolite yields. In the figure, αKG is denoted as Oxoglutarate and Ribose-5-Phosphate as R5P.

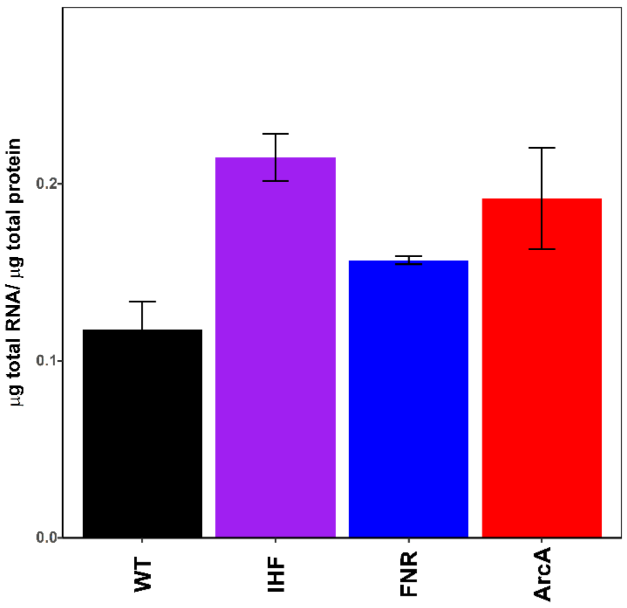


**Fig. S11.** Bar-plot depicting the experimentally derived total RNA/total protein ratio in all the strains (WT, Δ*ihf*, Δ*fnr* & Δ*arcA*) depicted as μg total RNA/ μg total RNA. Error bars represent standard deviations within replicate samples.

| STRAINS | GROWTH RATE ($h^{-1}$) | GLUCOSE UPTAKE RATE (mmol/gDCW/h) | YIELD OF ETHANOL (g/g glucose) | YIELD OF FORMATE (g/g glucose) | YIELD OF ACETATE (g/g glucose) | YIELD OF LACTATE (g/g glucose) | YIELD OF SUCCINATE (g/g glucose) | YIELD OF PYRUVATE (g/g glucose) | YIELD OF BIOMASS (g/g glucose) | AMMONIA UPTAKE RATE (mmol/gDCW/h) |
|---|---|---|---|---|---|---|---|---|---|---|
| WT | 0.38 ± 0.007 | 16.63 ± 0.5 | 0.18 ± 0.01 | 0.35 ± 0.01 | 0.21 ± 0.01 | 0.02 ± 0 | 0.09 ± 0.01 | 0.02 ± 0 | 0.13 ± 0 | 6.07 ± 0.5 |
| Δ*ihf* | 0.29 ± 0.007 | 13.26 ± 0.23 | 0.19 ± 0 | 0.36 ± 0.01 | 0.23 ± 0 | 0.04 ± 0.017 | 0.07 ± 0.03 | 0.003 ± 0 | 0.12 ± 0 | 4.76 ± 0.63 |
| Δ*fnr* | 0.29 ± 0.008 | 12.12 ± 0.62 | 0.18 ± 0.01 | 0.39 ± 0.02 | 0.23 ± 0 | 0.04 ± 0 | 0.05 ± 0 | 0.002 ± 0 | 0.13 ± 0.01 | 4.4 ± 0.53 |
| Δ*arcA* | 0.32 ± 0.015 | 13.99 ± 0.44 | 0.18 ± 0 | 0.33 ± 0.02 | 0.21 ± 0.01 | 0.01 ± 0 | 0.12 ± 0.01 | 0.002 ± 0 | 0.13 ± 0 | 4.99 ± 0.21 |

**Table S1:** Physiological characterization of the strains in anaerobic fermentation of glucose. The measurements of growth rate, glucose and ammonia uptake rate, and yields of mixed acid fermentation products were obtained from three biological replicates (n = 3). Yields were calculated by normalizing secretion rates with its glucose uptake rate. The errors indicate standard deviations within the replicates.

| **Strain Name** | **Genotype** | **Source** |
|---|---|---|
| *E. coli* K12 MG1655 WT | F-, *λ*-, *rph-1* | Keio collection<br>*CGSC #6300* |
| *E. coli* K12 MG1655 Δ*arcA* | F-, *λ*-, *rph-1, Δarca::kan* | This study |
| *E. coli* K12 MG1655 Δ*fnr* | F-, *λ*-, *rph-1*, Δ*fnr::kan* | This study |
| *E. coli* K12 MG1655 Δ*ihf* | F-, *λ*-, *rph-1*, Δ*ihfA::FRT*<br>Δ*ihfB::kan* | This study |

**Table S2.** The *E. coli* strains used in this study.

| Name | Primer Sequence (5'-3') |
| --- | --- |
| ArcA KO Fwd | CTTTTGTACTTCCTGTTTCGATTTAGTTGGCAATTTAGGTAGCAAACGTGTAGGCTGGAGCTGCTTCG |
| ArcA KO Rev | CGGCGCTAAAAAGCGCCGTTTTTTTTGACGGTGGTAAAGCCGACCGGGGATCCGTCGACC |
| FNR KO Fwd | CTAAAAAGATGTTAAAATTGACAAATATCAATTACGGCTTGAGCAGACCTGTGTAGGCTGGAGCTGCTTCG |
| FNR KO Rev | CAGAAAAATTTAATGATATGACAGAAGGATAGTGAGTTATGCGGAAAAACCGGGGATCCGTCGACC |
| IhfA KO Fwd | GAGGCATTAAAAGAGCGATTCCAGGCATCATTGAGGGATTGAACCTGTGTAGGCTGGAGCTGCTTCG |
| IhfA KO Rev | CAGTGAAAAGAAAAAAGGCCGCAGAGCGGCCTTTTTAGTTAGATCAGACCGGGGATCCGTCGACC |
| IhfB KO Fwd | GCAGCAACAGCAGCCGCTTAATTTGCCTTTAAGGAACCGGAGGAATCGTGTAGGCTGGAGCTGCTTCG |
| IhfB KO Rev | GCACCCGACAGGTGCTTTTCTCTCGTTCAAGTTTGAGTAAAAAACCCGGGGATCCGTCGACC |
| ArcA Det Fwd | TTTTGACACTGTCGGGTCCTGAGGGAAAGT |
| ArcA Det Rev | TTGGGAACCAGTGTGCTGGTGGTGG |
| FNR Det Fwd | CTGTAAACATTAAACAATTTGTGCCAGCTTG |
| FNR Det Rev | CGTCCTGGTTAGGATCGATAACAACGA |
| IhfA Det Fwd | CGTAAATCAGGTAGTTGGCGTAAACTTATTTGACG |
| IhfA Det Rev | TTAAGAGAAGCGCCAGCAGCATCAAA |
| IhfB Det Fwd | ACGCAATGGCTGAAGCTTTCAAAGCA |
| IhfB Det Rev | TCCCTGTTTATGGAAAGTGTGCAACTTTGT |

**Table S3.** Primer designs used in this study

| **Comparison** | Δ*fnr* compared to WT | | Δ*ihf* compared to WT | | | Δ*arcA* compared to WT | | |
|---|---|---|---|---|---|---|---|---|
| **Gene** | **RT-PCR (FC ± Std dev)** | **RNA-Seq (FC)** | **Gene** | **RT-PCR (FC ± Std dev)** | **RNA-Seq (FC)** | **Gene** | **RT-PCR (FC ± Std dev)** | **RNA-Seq (FC)** |
| *ompF* | 0.145592 ± 0.05 | 0.090873 | *hdeA* | 0.008088 ± 0.003 | 0.00897421 | *appC* | 0.161544 ± 0.037 | 0.150726 |
| *ilvE* | 0.31864 ± 0.028 | 0.258816 | *alaE* | 0.023683 ± 0.002 | 0.01923663 | *asnA* | 0.156041 ± 0.076 | 0.129408 |
| *nrdF* | 0.450625 ± 0.081 | 0.125 | *ilvE* | 0.101532 ± 0.009 | 0.09473229 | *feoB* | 0.213159 ± 0.095 | 0.189465 |
| *ackA* | 0.4 ± 0.015 | 0.464997 | *aceB* | 0.101532 ± 0.025 | 0.11502346 | *oppF* | 0.185565 ± 0.052 | 0.176777 |
| *Acs* | 9.781122 ± 0.773 | 7.012846 | *glnA* | 5.35171 ± 0.458 | 5.434917 | *sucA* | 5.656854 ± 0.958 | 10.19649 |
| *sdhA* | 9.565 ± 0.864 | 9.57983 | *hsdR* | 10.55606 ± 1.001 | 8.446723 | *sdhA* | 17.38776 ± 1.525 | 17.87659 |
| *phoH* | 5.578975 ± 0.801 | 5.278032 | *ompF* | 7.674113 ± 1.222 | 7.014112 | *glcD* | 48.50293 ± 3.915 | 135.0326 |
| *ppsA* | 2 ± 0.275 | 2.143827 | *glnK* | 4.594793 ± 0.429 | 16.98619 | *acs* | 12.12573 ± 1.885 | 50.91433 |

**Table S4:** Validation of RNA-seq using qRT-PCR. Measurements were performed on biological duplicates and technical triplicates (n=6).

| Name | RT-PCR Primer Sequence (5'-3') | Name | RT-PCR Primer Sequence (5'-3') |
|---|---|---|---|
| ompF Fwd | AGGCTTTGGTATCGTTGGTG | aceB Fwd | CTGCGTGACCATATTGTTGG |
| ompF Rev | TTGTTCGCGTCGTACTTCAG | aceB Rev | CATCGTCACTGCCTGTCTGT |
| ilvE Fwd | TTTCGCAGAGCATTGATGAG | glnA Fwd | ACCCGCTTCAATACCATGAC |
| ilvE Rev | TTACTCCCATGCCAACATCA | glnA Rev | AGCCGTTATCACCGAACATC |
| nrdF Fwd | CACTCACGCCTCATGAAGAA | hsdR Fwd | GATCCAGTCCCAGCGTTTTA |
| nrdF Rev | GGCATCGACATCTTTGGTCT | hsdR Rev | TCCGTCAGCCCTTTAATGTC |
| ackA Fwd | TACCTCTACGCCCTGCCTTA | glnK Fwd | CGTCACCGAAGTGAAAGGTT |
| ackA Rev | CCGGTTTGTTCAGCATTTTT | glnK Rev | AGCAATCGCCACATCAATTT |
| acs Fwd | GGGTGACCGGACACAGTTAC | appC Fwd | GATCCTCGCCACTCACTCAT |
| acs Rev | TTGACCTGATGCTTGTCCAC | appC Rev | ATTAACAGCCAGGCCATACG |
| rpoB Fwd | ACTCAGCTGACCCCAGAAGA | asnA Fwd | CACTTCGTACACAGCCAGGA |
| rpoB Rev | ACCGGATACACCGTTTGGTA | asnA Rev | CCAATCCCGACAAGGAATAC |
| sdhA Fwd | CGTACTGGTCACCGAAGGTT | feoB Fwd | TGCCGGTCTATCATGTACCA |
| sdhA Rev | CCTTCACGGATTTCGATCAT | feoB Rev | TTGAAAGCGCTCAGGAAAAT |
| phoH Fwd | GCGCAATTGCACTATCTGAA | oppF Fwd | AAGGCGATGATGCTGAAAGT |
| phoH Rev | GGGTGACGATAATCCTGTCG | oppF Rev | AGAATAAGAGCACGCGCAAT |
| ppsA Fwd | ACTGGGAACCGATCATGAAG | sucA Fwd | CCCGTCTGGACAGACTTGAT |
| ppsA Rev | TTCTGTTGCATCTCCACAGC | sucA Rev | CGACATGTTCAGGGTTTCCT |
| hdeA Fwd | GAAGATTTCCTGGCTGTGGA | glcD Fwd | GCTGGATTCACCTGGTTTTG |
| hdeA Rev | ACGGTTGCAATACCCTGAAC | glcD Rev | CCGAGTCAAAGCTGGCTAAC |
| alaE Fwd | GTAGCGATTCCGGTGAACAT | | |
| alaE Rev | CAGGATATCCGCCAGATTTT | | |

**Table S5.** qRT-PCR primer designs used in this study
